# DAF-18/PTEN inhibits germline zygotic gene activation during primordial germ cell quiescence

**DOI:** 10.1101/2020.09.27.314948

**Authors:** Amanda L. Fry, Amy Webster, Rojin Chitrakar, L. Ryan Baugh, E. Jane Albert Hubbard

## Abstract

Quiescence, an actively-maintained reversible state of cell cycle arrest, is not well understood. PTEN is one of the most frequently lost tumor suppressors in human cancers and regulates quiescence of stem cells and cancer cells. In *C. elegans* mutant for *daf-18*, the sole *C. elegans* PTEN ortholog, primordial germ cells (PGCs) divide inappropriately in starvation conditions, in a TOR-dependent manner. Here, we further investigated the role of *daf-18* in maintaining PGC quiescence. We found that maternal or zygotic *daf-18* is sufficient to maintain cell cycle quiescence, that *daf-18* acts in the germ line and soma, and that *daf-18* affects timing of PGC divisions in fed animals. Importantly, our results also implicate *daf-18* in zygotic germline gene activation, though not in germline fate specification. However, TOR is less important to zygotic germline gene expression, suggesting that in the absence of food *daf-18*/PTEN prevents inappropriate germline zygotic gene activation and cell division by distinct mechanisms.

## Introduction

Cell cycle quiescence is an actively maintained state of non-proliferation. The best characterized quiescent state known as “G_0_,”is associated with exit from the cell cycle after M phase. Depending on the cell type, arrested cells adopt a variety of altered metabolic states while they await an activation cue, usually a growth factor signal (Cheung and Rando 2013; Coller 2019a; Shyh-Chang et al. 2013). Response to these cues ultimately leads to post-transcriptional modifications of cell cycle proteins, returning the cell to a cycle of continuous G1, S, G2 and M phases (Coller 2019b).

Cell cycle quiescence can also occur in the G2 phase of the cell cycle. Though far less well-understood, G2 quiescence has been observed in vertebrate muscle stem cells, *Drosophila* neural stem cells, and *C. elegans* germ cells (Fukuyama et al. 2006; Nguyen et al. 2017; Otsuki and Brand 2018). Proper regulation of quiescence is particularly important in founder cells such as stem cells (van Velthoven and Rando 2019). Inappropriate quiescence of these cells can cause loss of downstream differentiated cell populations, aberrant tissue homeostasis, and failure to repair tissue damage. Inappropriate exit of these cells from quiescence can prematurely deplete the stem cell pool and similarly cause defects in tissue homeostasis. Thus, understanding how quiescence is modulated is crucial for understanding the behavior of stem cells *in vivo*. In addition, this understanding is important for the development of new therapies to target cancer stem-like cells since quiescent cancer stem cells are refractory to common chemotherapies and are likely responsible for recurrent cancer (Batlle and Clevers 2017; Cheung and Rando 2013; Cho et al. 2019; Pavlova and Thompson 2016).

The regulation of both *Drosophila* neural stem cell and *C. elegans* germ cell quiescence has been linked to organismal nutrient status by insulin-PI3-kinase (PI3K) signaling and PTEN. <u>P</u>hosphatase and <u>Ten</u>sin homolog (PTEN) regulates nutrient-sensitive pathways and inhibits proliferation of stem/progenitor cells (Hill and Wu 2009; Sun et al. 1999; Kimura et al. 2003; Yilmaz et al. 2006; Zhang et al. 2006; White et al. 2014), cancer cells (Worby and Dixon 2014), and cancer stem cell-like populations (Duan et al. 2015). PTEN is the second-most commonly lost tumor suppressor in human cancers (Hill and Wu 2009; Leslie and Downes 2004). Molecularly, PTEN functions as a lipid and protein phosphatase (Worby and Dixon 2014). It dephosphorylates the lipid second messenger phosphatidylinositol _(3,4,5)_ tri-phosphate (PIP_3_), converting it to PI_(4,5)_P (PIP_2_). Thus, PTEN opposes the activity of PI3K, diminishing the effects of upstream receptors (such as growth factor receptors and the insulin receptor) on proteins activated by PIP_3_, such as AKT, a positive effector of cell cycle progression (Keniry and Parsons 2008). Interestingly, PTEN loss in mammals can render tumor cells insensitive to dietary restriction (Kalaany and Sabatini 2009; Curry et al. 2013), consistent with PTEN mediating nutritional control of cell proliferation. The breadth of mechanisms by which PTEN regulates cell quiescence is not fully understood. Understanding the mechanisms by which PTEN regulates G2 stem cell quiescence in the context of normal development may also uncover additional roles important for cancer.

The sole PTEN ortholog in *C. elegans* is DAF-18, and it was the first PTEN homolog to be described in a genetically tractable model organism. It was identified as a negative regulator of insulin signaling in dauer formation (dauer is a stress-resistant larval stage; “daf” stands for “abnormal <u>DA</u>uer <u>F</u>ormation”) (Riddle et al. 1981; Vowels and Thomas 1992; Ogg and Ruvkun 1998). In *C. elegans* the two primordial germ cells (PGCs) are born relatively early in embryogenesis but do not divide again until worms have hatched and begin feeding. Loss of *daf-18* causes G2-arrested PGCs to proliferate inappropriately in first larval stage worms (L1) despite starvation (Fukuyama et al. 2006). Loss of *daf-18* also interferes with later germ cell cycle arrest during dauer (Narbonne and Roy 2006) via non-autonomous activity in the somatic gonad (Tenen and Greenwald 2019). Remarkably, human PTEN can rescue *daf-18* mutant dauer and longevity phenotypes (Solari et al. 2005), indicating conserved function.

The current model for regulation of PGC cell cycle by DAF-18 in starved L1 worms is that DAF-18 regulation occurs downstream of insulin signaling, through PI3K, Akt and TOR, but is independent of DAF-16 FOXO (Baugh and Sternberg 2006; Fukuyama et al. 2015; Fukuyama et al. 2006; Fukuyama et al. 2012; Kasuga et al. 2013; Zheng et al. 2018). While *C. elegans* germ cells arrest in the G2 phase of the cell cycle during L1 starvation (Fukuyama et al. 2006), most somatic cells arrest in G1 in a *daf-16*-dependent fashion through regulation of the cyclin dependent kinase inhibitor CKI-1 (Baugh and Sternberg 2006; Hong et al. 1998).

To further understand how DAF-18 PTEN acts to maintain L1 PGC quiescence, we further characterized the role of DAF-18, taking advantage of a null allele *daf-18(ok480)* and a live marker for germ cells. Our results suggest that DAF-18 influences the onset of germ cell division in both starved and fed L1 larvae, and that DAF-18 appears to act in both the germ line and the soma to regulate PGC quiescence. We also determined that either maternal or zygotically supplied DAF-18 can maintain quiescence. Finally, we investigated the relationship between germline zygotic gene activation and PGC quiescence. We discovered that transcriptional quiescence accompanies cell division quiescence, and that DAF-18 maintains both. However, we found that transcriptional quiescence is not as sensitive to TOR as is cell cycle quiescence, suggesting that a non-PI3K-dependent activity of DAF-18 PTEN may underlie transcriptional quiescence.

## Results

### DAF-18 influences the timing of PGC division onset in both starved and fed L1 larvae

The primordial germ cells (PGCs) Z2 and Z3 are born by division of the P4 cell in the embryo and do not divide further until L1 larvae have hatched and begin feeding. In worms bearing loss-of-function mutations in *daf-18*, PGCs divide in L1 larvae in the absence of food (Fukuyama et al. 2006). To determine when the PGCs of starved *daf-18* mutants first divide relative to fed controls, we performed a time-course analysis in worms carrying *naSi2*, a germline-expressed transgene of mCherry fused to histone H2B (Roy et al. 2018) (Figure 1A). We first established that *daf-18* mutant embryos collected at the 2-4 cell stage hatch at the same time as wild-type embryos (Figure S1). We then monitored PGC divisions in L1 larvae that had been tightly synchronized relative to hatching (see Methods) in wild-type and *daf-18* mutants in the absence and presence of food. We found that PGCs in starved *daf-18* mutants began to divide at the same time (4 hours post-collection; see Methods) as did fed wild-type worms (Figure 1B,C). Surprisingly, fed *daf-18* mutant PGCs also divide prematurely compared to fed wild-type L1 larvae (Figure 1C). Thus, PGCs divide several hours earlier in fed *daf-18* mutants than starved *daf-18* mutants. These results indicate that while *daf-18* mutants show food-independent PGC division that mimics the timing of PGC division in fed wild-type worms, PGCs in *daf-18* mutants also appear primed to divide such that they initiate divisions early in the presence of food.

**Figure 1.**
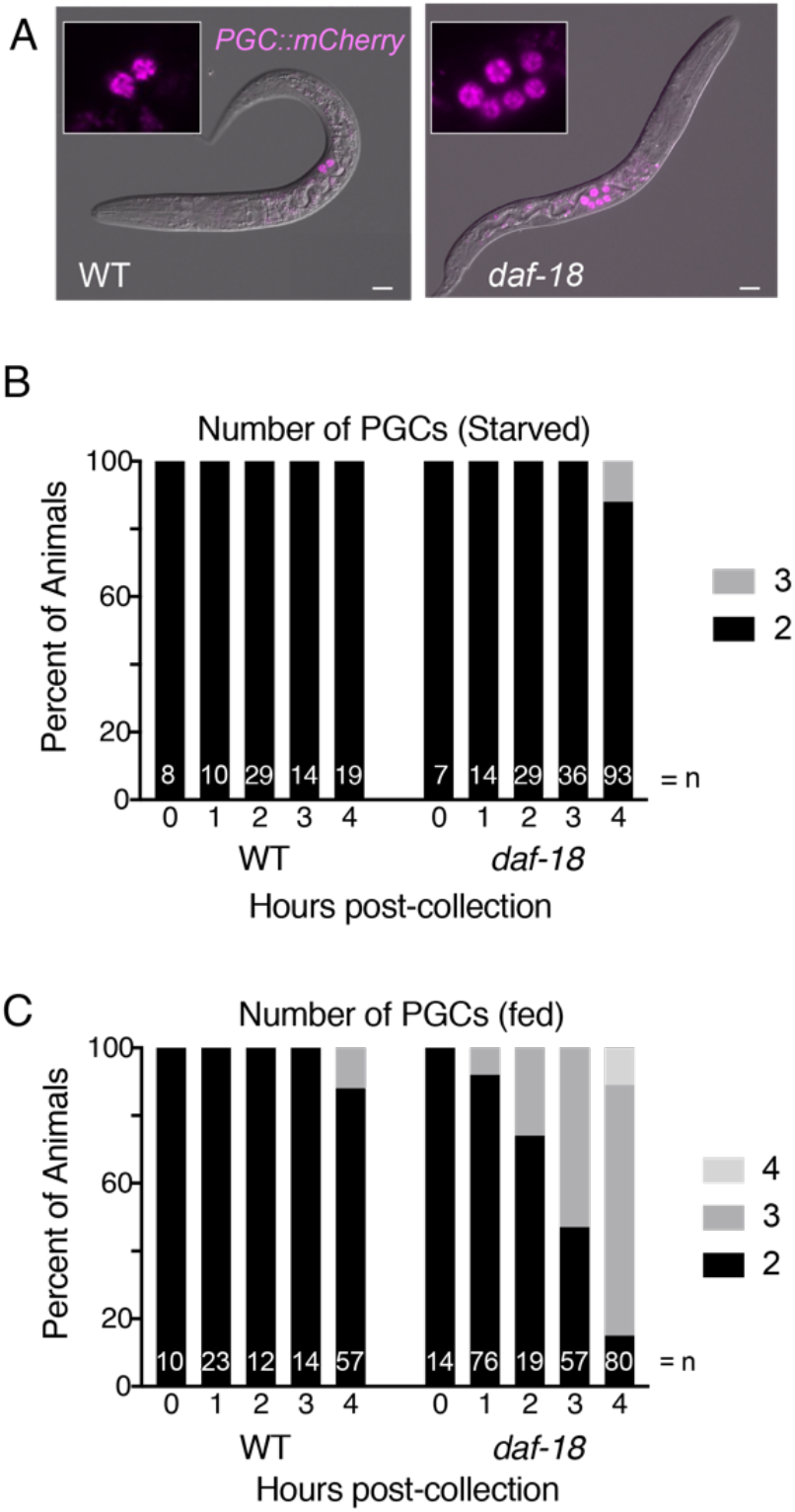
DAF-18 promotes PGC quiescence in both fed and starved larvae. **A**. Wild type (WT) and *daf-18(ok480)* mutant L1 larvae, starved for 3 days after hatching, show 2 or an average of 6 primordial germ cells (PGCs), respectively. The marker *PGC::mCherry* (*naSi2*) is expressed in the PGCs (and in ∼10% of animals, expression is seen in one additional cell in the head). Scale bar represents 10μm. **B. and C**., Animals were synchronized within 2 hours of hatching and monitored for PGC divisions during starvation (**B**) and in the presence of food (**C**). Summary of 2 replicate experiments.

### DAF-18 appears to act in both the soma and germ line to regulate PGC quiescence

Previous studies found that transgenic expression of *daf-18(+)* genomic sequences restores PGC quiescence (Fukuyama et al. 2012). These transgenes were expressed from extrachromosomal arrays, a technique which often precludes germline expression (Kelly and Fire 1998). We generated a similar transgene bearing the *daf-18(+)* genomic sequence on an extrachromosomal array, with a trans-splice to GFP::H2B to follow expression without altering with the DAF-18 protein itself (see Methods). We confirmed that PGC quiescence was fully restored in animals bearing this *daf-18(+)* transgene (Figure 2, “genomic array”). Consistent with other studies showing expression of *daf-18* reporters in varied tissues (Brisbin et al. 2009; Masse et al. 2005; Nakdimon et al. 2012; Packer et al. 2019), we observed widespread expression of GFP, including within the intestine, neurons, and hypodermis. We did not observe pression from our *daf-18(+)* genomic transgene in the PGCs (Figure S2).

**Figure 2.**
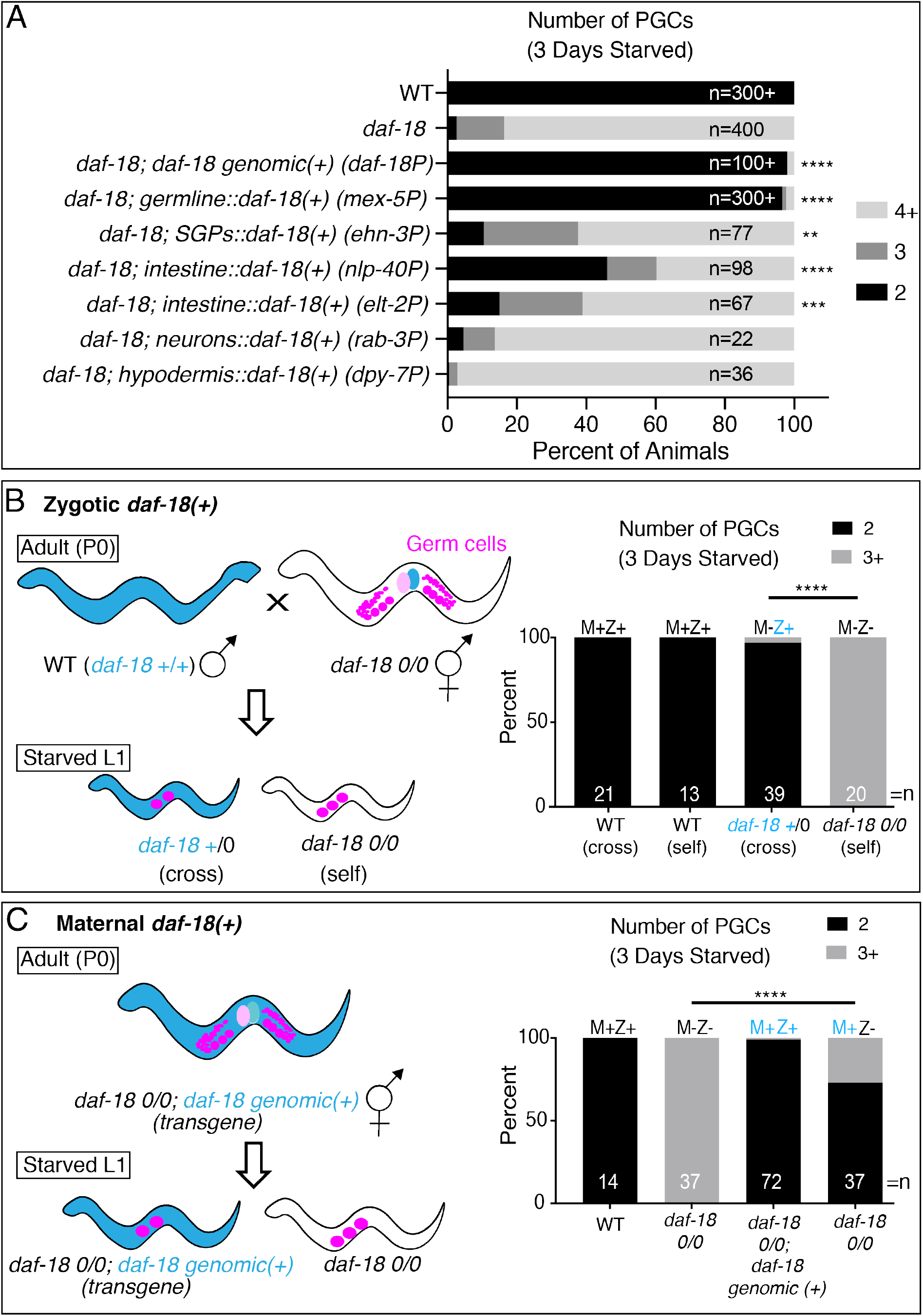
DAF-18 acts in the germ line and soma to suppress PGC division during starvation. **A**. The number of PGCs in L1s was assessed after 3 days of starvation. Genotypes indicate the promoters used to drive tissue-restricted expression of the *daf-18* coding region containing all introns and *daf-18* 3’UTR, with the exception of *germline::daf-18(+)*, which contains the same *daf-18* coding region with introns, but the *nos-2* 3’UTR. All animals carry *PGC::mCherry*, except for *daf-18; germline::daf-18(+)*, which carries the PGC marker *glh-1::GFP* or no additional PGC marker (*germline::daf-18* is tagged with GFP; see Figure S2). n values display the total number of animals examined for each genotype. Chi-square statistical tests (or two-sided Fisher’s exact test for *ehn-3P, elt-2P, rab-3P and dpy-7P*) were performed on each genotype compared to its own experimental control group (*daf-18(ok480)* mutant with *PGC::mCherry*) and significance is displayed: **p<0.01, ***p<0.001, ****p<0.0001. Additional independently-generated transgenic lines were examined for several transgenes, with similar results found: 3 additional lines of *daf-18P::daf-18(+)* (suppressed *daf-18*), 2 additional lines of *nlp-40P::daf-18(+)* (suppressed *daf-18*), 1 additional line of *dpy-7P::daf-18(+)* (no suppression), and 1 additional line of *rab-3P::daf-18(+)* (no suppression). **B**. Zygotic DAF-18 maintains PGC quiescence. Blue color indicates the presence of *daf-18(+)*. Strategy is shown to generate Maternal-Zygotic+ (M-Z+) by crossing paternal *daf-18(+)* to *daf-18* mutant hermaphrodites. This strategy gives starved L1 cross-progeny one wild-type copy of *daf-18* in the absence of maternal *daf-18*. Cross-versus self-progeny were identified by the presence/absence of a bright somatic fluorescent marker from the paternal P0 strain (carrying *cdc-42::GFP*). Number of PGCs were assessed after 3 days of starvation of L1 progeny. Statistical significance determined by two-sided Fisher’s exact test. ****p<0.0001. **C**. Maternal DAF-18 also maintains PGC quiescence. Blue color indicates the presence of *daf-18(+)*. Strategy is shown to provide maternal *daf-18(+)* in the presence (M+Z+) or absence (M+Z-) of zygotic *daf-18(+)* by analyzing the progeny of *daf-18* mutants (0/0) carrying a *daf-18(+)* genomic transgene. This transgene was stochastically lost in progeny, yielding M+Z+ or M+Z-L1 larvae, which were assessed for PGC numbers after 3 days of starvation. Statistical significance determined by two-sided Fisher’s exact test. ****p<0.0001.

To determine whether *daf-18(+)* expression from specific somatic tissues might also restore PGC quiescence, we generated plasmids driving *daf-18(+)* expression (similar to genomic array above, including all introns and trans-spliced GFP::H2B, but with alternative sequences 5’ to the *daf-18* ATG) from various tissue-restricted promoters. We found that *daf-18(+)* expression in the intestine or somatic gonad precursors, but not in the hypodermis or neurons, partially restored PGC quiescence in *daf-18* mutants, albeit not to the same level as the genomic array (Figure 2A).

Since *daf-18* is reported to be highly expressed in the germ cells (Brisbin et al. 2009; Packer et al. 2019; Suzuki and Han 2006), and regulates L1 PGC division (Fukuyama et al. 2006), we tested whether germline-expressed *daf-18(+)* can support PGC quiescence in the absence of food. We expressed *daf-18(+)* in the PGCs using a transgene bearing a construct consisting of a germline promoter (*mex-5P*) and 3’ UTR (*nos-2*) flanking the *daf-18* coding region, trans-spliced to GFP (fused to a plextrin homology (PH) domain, which enhanced membrane localization). DNA bearing these sequences was inserted into the genome using CRISPR/Cas9 (Dickinson et al. 2013). These regulatory regions limit expression to the germ line, as seen with our PGC::mCherry marker using the same regulatory regions (Figure S2A). This transgene was highly expressed in the PGCs and, in contrast to partial rescue from *daf-18(+)* driven from intestine and somatic gonad precursor promoters, it nearly fully restored PGC quiescence to *daf-18* mutants (Figure 2A).

Taken together, our results suggest that germline DAF-18 plays a major role in regulating PGC quiescence, but that DAF-18 in somatic tissues (including intestine and somatic gonad precursors) may contribute to PGC quiescence. GFP expression was observed from the genomic and all somatic transgenes in the expected tissues and none was observed in the germ line (Figure S2). None of the genes from which we chose regulatory regions for somatic tissue-restriction drive high levels of transcription (>100 transcripts per million; see Figure S2) in the germ line based on single cell RNA sequencing (Cao et al. 2017; Packer et al. 2019). Nevertheless, it remains formally possible that somatic promoter arrays express *daf-18(+)* in the germ line below the level of detection and that this could contribute to the rescue observed with these transgenes.

### Maternal or zygotic daf-18 activity maintains PGC quiescence in starved L1 larvae

PGC divisions normally occur early in larval development, a time when maternal or zygotic *daf-18* could regulate PGC divisions. To determine whether zygotic *daf-18(+)* alone is sufficient to confer quiescence, we crossed *daf-18(0)* mutant hermaphrodites with *daf-18(+)* males to generate *daf-18* heterozygous progeny lacking maternal *daf-18(+)* but expressing zygotic *daf-18(+)* (M-Z+) and inspected them for PGC quiescence during L1 starvation. We found that PGC quiescence was restored, indicating that zygotic *daf-18(+)* is sufficient for normal PGC quiescence (Figure 2B).

We then tested whether maternal *daf-18(+)* alone could also maintain PGC quiescence. We examined progeny from *daf-18(0)* mothers carrying a *daf-18(+)* genomic rescuing array (described above). Progeny that had lost the *daf-18(+)* array (see Methods) nevertheless showed PGCs arrested appropriately during starvation (Figure 2C), suggesting that maternal *daf-18(+)* is also sufficient for PGC quiescence.

### Marks of PGC transcriptional activation are elevated in starved daf-18 mutants prior to cell division

Several marks of active transcription in the PGCs are thought to be dependent on exposure to food (H3K4me2, H3K4me3, and “active” Pol II). The levels of these marks are reported to be low or absent in PGCs of late-stage embryos or starved L1 larvae relative to PGCs of fed L1 larvae or relative to somatic cells (Butuci et al. 2015; Checchi and Kelly 2006; Furuhashi et al. 2010; Schaner et al. 2003). Specifically, the levels of H3K4me2 detection in PGCs of L1 larvae is low relative to somatic cells prior to feeding and is elevated after feeding and prior to cell division (Schaner et al. 2003). In addition, “active” Pol II (phosphorylated Ser2 on the C terminal domain of Pol II; P-Ser2) is more readily detected in germ cells once L1 larvae begin to feed (Butuci et al. 2015).

To determine whether these molecular marks are elevated in PGCs of *daf-18* mutants in the absence of food, we examined levels of antibodies detecting H3K4me2, H3K4me3, and P-Ser2 of RNA Pol II in starved *daf-18(+)* and *daf-18(0)* relative to somatic cells. We found that while marks of H3K4me2 and H3K4me3 were detectable in the PGCs of starved wild-type worms, their levels were significantly lower relative to the surrounding somatic cells (Figure 3). In contrast, PGCs of starved *daf-18* mutants shortly after hatching and prior to cell division, displayed elevated levels of H3K4me2 and H3K4me3, comparable to the surrounding somatic cells (Figure 3B,C; Figure S3). This observation suggests that the chromatin in *daf-18* mutant PGCs may be permissive for transcription during starvation.

**Figure 3.**
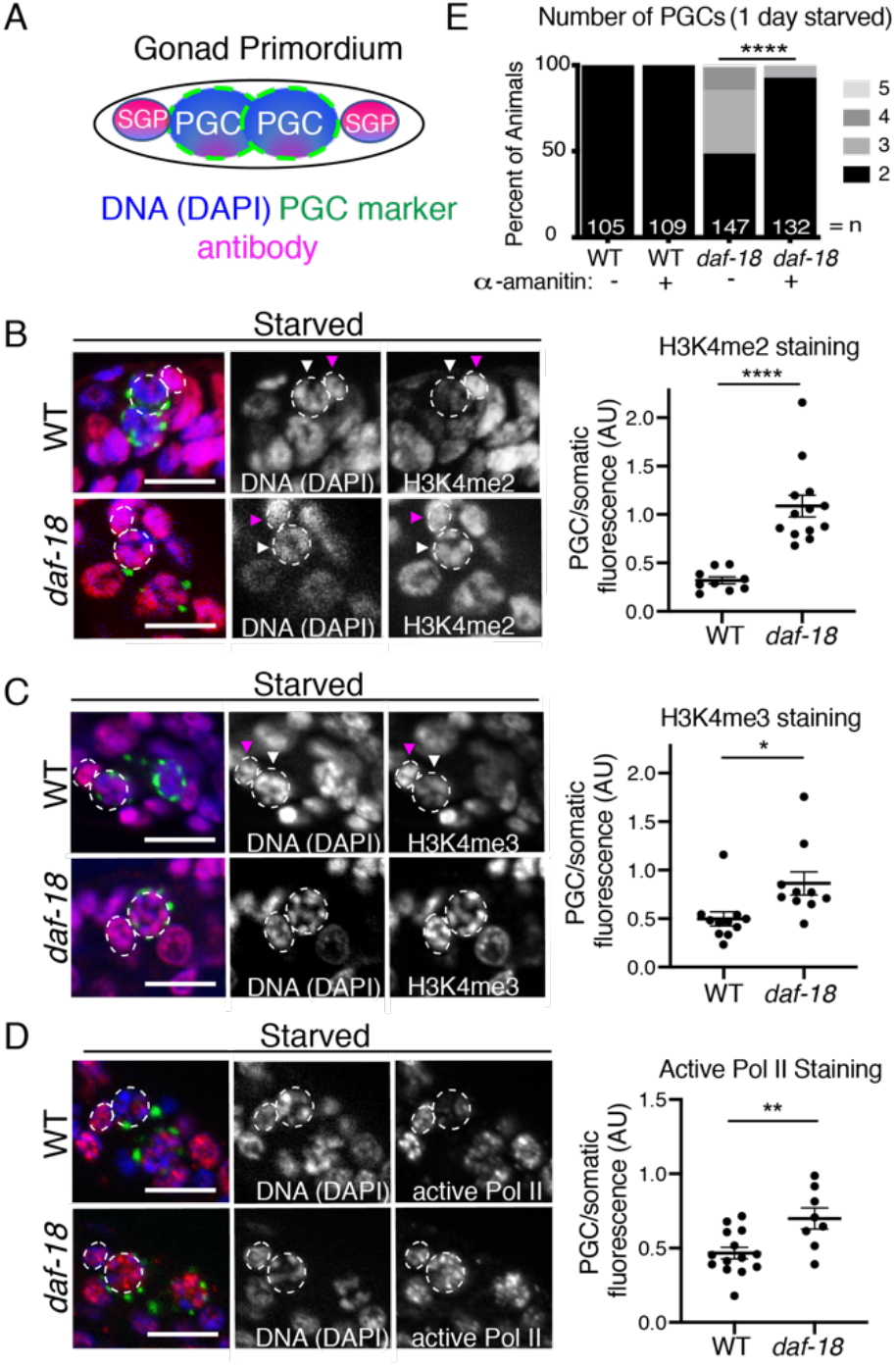
Marks of active transcription are inappropriately elevated in the PGCs of starved *daf-18* mutants. **A**. Schematic of gonad primordium, containing 2 PGCs, each with 1 neighboring somatic gonad precursor (SGP), and the stains/marker seen in each tricolor image (left panels of B-D). PGC marker is *glh-1::GFP*. **B**. Antibodies CMA303 (H3K4me2), **C**. ab8580 (H3K4me3), and **D**. H5 (Pol II P-ser2) were used to stain whole starved L1s. **B-D**. To illustrate PGC versus somatic fluorescence intensity, images in B and C show Z-projections of image stacks taken through the PGCs, while D shows single slices (Apotome). Tricolor images (left panels show DNA (DAPI/blue), immunofluorescence (magenta), and a PGC marker (encoded perinuclear *glh-1::GFP*). All animals were starved up to 2 hours (synchronized relative to hatching). Dashed circle indicates one PGC in each image, and dashed smaller ellipse indicates its nearest somatic cell (likely its SGP). White arrowheads point to PGCs, and pink arrowheads point to neighboring somatic cells. Mean staining fluorescence was quantified for each PGC nucleus and its nearest somatic cell nucleus (from single image slices), and is displayed in the graph as a ratio. Each dot in the graph is an average value per worm (2 PGC/SGP values averaged). Statistical significance determined by two-tailed T-test. *p<0.05, **p<0.01, ****p<0.0001. **E**. Transcription by Pol II was inhibited with alpha-amanitin (10μg/mL) and PGC numbers were assessed after 1 day (∼24h) of starvation. Statistical significance determined by two-sided Fisher’s exact test. ****p<0.0001. All scale bars represent 10μm.

To assess whether transcription is occurring in PGCs of starved *daf-18* mutant L1 larvae, we examined the levels of P-Ser2. We detected low levels in PGCs of starved wild-type worms compared to surrounding somatic cells. Similar to what we observed with active chromatin marks, P-Ser2 is markedly elevated in PGCs in starved *daf-18* mutant L1 larvae, reaching levels roughly equivalent to nearby somatic cells (Figure 3D).

### Inhibiting transcription suppresses PGC divisions in starved daf-18 mutants

Since we observed that PGCs in starved *daf-18* mutants display inappropriately high levels of marks of transcriptional activation shortly after hatching and well before the time that PGCs would divide, we hypothesized that inhibiting transcription would block inappropriate PGC divisions. To test this hypothesis, we inhibited Pol II using *α*-amanitin (Chafin et al. 1995; Rudd and Luse 1996). We allowed *daf-18* mutant and wild-type embryos to hatch in buffer with no food, with or without *α*-amanitin. This treatment significantly reduced the proportion of starved *daf-18* mutant animals with ≥3 PGCs after one day of L1 starvation (Figure 3E). Taken together with the elevated marks of active transcription, these results suggest that activation of PGC transcription precedes inappropriate PGC divisions in starved *daf-18* mutants and may be required for such divisions.

### Levels of zygotic *VBH-1* and *GLH-1* are inappropriately elevated in PGCs of starved daf-18 mutants prior to cell division

It was previously shown by mRNA-FISH that the transcription of genes encoding five different P granule associated proteins is induced in PGCs upon L1 feeding (Wong et al. 2018). To assess this germline expression in live worms, we tagged one of them, VBH-1 (<u>V</u>asa and <u>b</u>elle-like RNA <u>h</u>elicase) with GFP using CRISPR/Cas9. To assess zygotic expression of the GFP::VBH-1 protein fusion, we crossed it in from the male, creating (M-Z+) F1 progeny carrying one paternal copy of GFP::VBH-1. Since these F1 progeny come from non-GFP mothers, any green fluorescence is the result of zygotic expression (Figure 4A). In the wild-type genetic background, zygotic perinuclear GFP::VBH-1 was only barely detectable in starved L1 PGCs. In contrast, PGCs in starved *daf-18* mutants displayed bright zygotic GFP::VBH-1. The intensity of this perinuclear fluorescence is equivalent to that of fed wild-type animals, and is observed within 1 hour of hatching, several hours before the PGCs begin dividing. In wild-type animals, after 4-5 hours of feeding, strong perinuclear expression was observed in the PGCs (Figure 4B).

**Figure 4.**
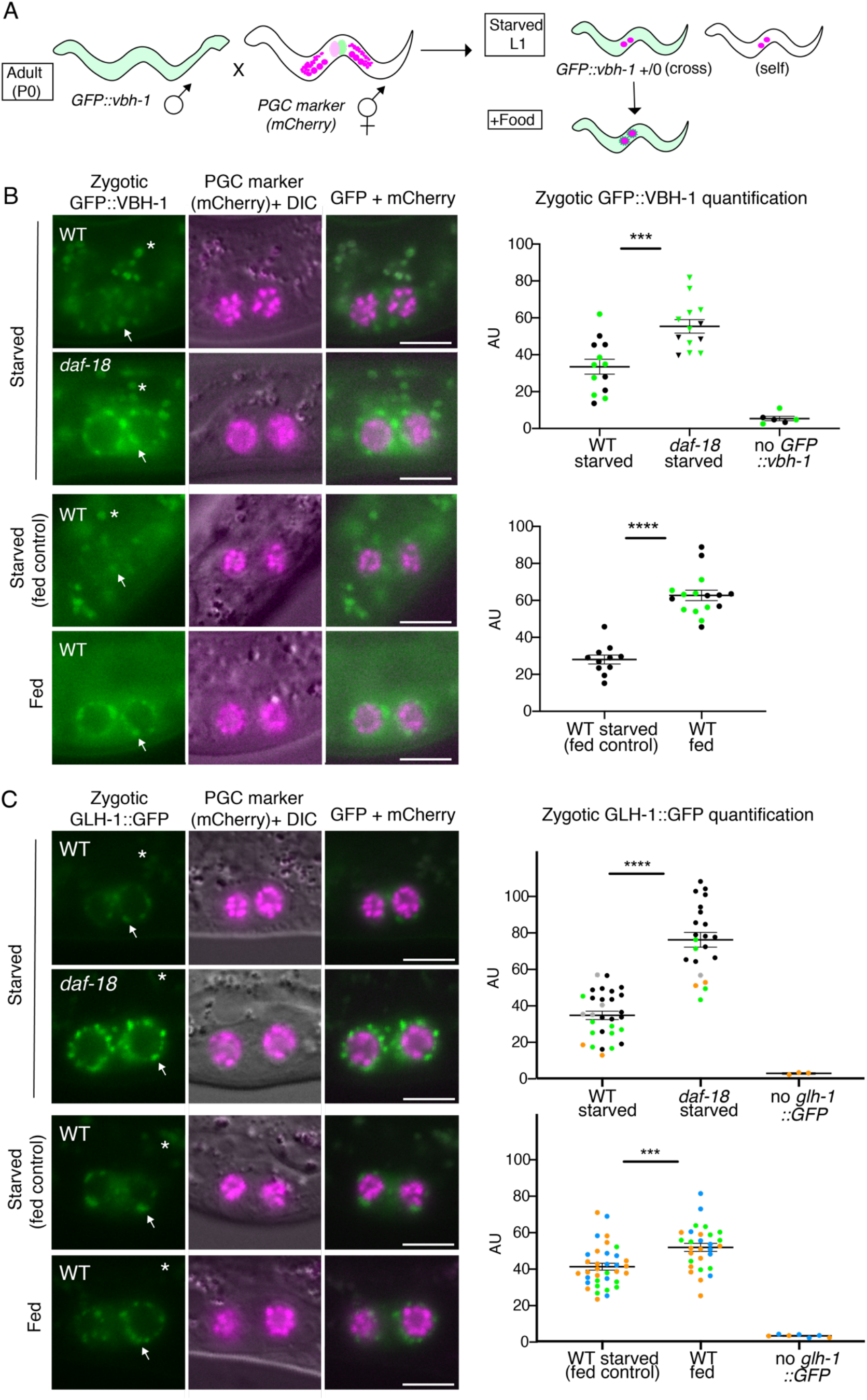
Levels of zygotic VBH-1 and GLH-1 are inappropriately elevated in PGCs of starved *daf-18* mutants. **A**. Schematic of strategy to observe strictly zygotic expression of germline genes *vbh-1* and *glh-1*, tagged with GFP. Single-copy GFP inserted into endogenous *vbh-1* or *glh-1* loci was crossed from the male into PGC::mCherry hermaphrodites. Crosses were performed in either the wild type (WT) or *daf-18(ok480)* mutant background. “Starved” L1 progeny were imaged up to 5 hours after hatching (with no food). “Starved (fed control)” progeny were imaged up to 24 hours after clean embryo preparation (no food), while “fed” L1 progeny were imaged at the same time, but upon 5 hours of feeding (after up to 19 hours post-embryo preparation). **B**. Representative images taken within 1 hour of hatching. Arrows point to zygotic perinuclear GFP::VBH-1 in PGCs. Exposure time is longer in B than in C, so gut granules (green auto-fluorescent dots, marked with asterisks and visible in DIC) are more prominent. Graphs show the results of two experiments pooled together (dots color-coded per experiment). Similar results for WT vs. *daf-18* starved were observed in an additional experiment on L1 larvae starved up to 24h (data not shown). **C**. Representative images taken up to 5 hours after hatching. Graphs show the results of 5 experiments (dots color-coded per experiment). Arrows point to zygotic perinuclear GFP::VBH-1 in PGCs. Gut granules (green auto-fluorescent dots) are marked with asterisks and visible in DIC. **B and C**. Images show epifluorescence. Mean perinuclear GFP fluorescence intensity (AU=Arbitrary Units) values are plotted to the right of each set of images. Each dot represents one animal (either average of 2 PGC values, or one PGC per animal). Non-cross progeny were identified by the lack of somatic GFP (“no GFP::vbh-1”) or lack of any GFP P-granules (“no GLH-1::GFP”; this transgene is much brighter and is clearly visible in starved WT PGCs). Mean +/-SEM. Statistical significance determined by two-tailed T-test. ***p<0.001 ****p<0.0001. All scale bars represent 10μm.

Since VBH-1 is broadly expressed at a low level in most somatic tissues in addition to its high expression in the germ line (Cao et al. 2017; Packer et al. 2019), we wondered whether *daf-18* mutants also zygotically express inappropriately high levels of GFP::VBH-1 in the soma. We observed faint, diffuse cytoplasmic expression of zygotic GFP::VBH-1 in somatic cells of the wild-type L1, and particularly in head neurons. However, we did not observe significant differences in this head expression between starved *daf-18* mutants and the wild type (Figure S4A).

To determine whether the inappropriate zygotic germline expression seen in starved *daf-18* mutant PGCs was specific to VBH-1, we investigated another germline gene reporter: GLH-1::GFP (<u>G</u>erm <u>L</u>ine <u>H</u>elicase), a component of germline-specific P granules (Gruidl et al. 1996). We used an available GLH-1::GFP CRISPR allele (Andralojc et al. 2017), and crossed it in from the male to examine zygotic protein expression. Like GFP::VBH-1, we found that in PGCs of starved *daf-18* mutant animals, the level of zygotic GLH-1::GFP expression was significantly elevated compared to the wild type within a few hours of hatching (Figure 4C).

In addition, we observed several differences between the zygotic germline activation of VBH-1 and GLH-1. GFP::VBH-1 is barely detectable in PGCs prior to feeding in the wild type and feeding strongly elevated expression relative to the starved condition (Figure 4). By contrast, the zygotic germline expression of GLH-1::GFP was relatively high in starved wild-type L1 worms. This is not completely surprising, since a previous report suggests that *glh-1* is transcribed zygotically in the embryo (Spencer et al. 2011). Nevertheless, zygotic VBH-1 levels are markedly elevated upon feeding. We also found that while the levels of expression of VBH-1 in PGCs of starved *daf-18* mutants are comparable to fed wild-type larvae, GLH-1 is expressed at a higher level in starved *daf-18* mutants than in the fed wild type. This observation suggests that *daf-18(+)* may contribute to a food-independent effect on zygotic germline gene expression.

We reasoned that if *daf-18* mutants elevate transcription of all germline genes, they should also express inappropriately high levels of our single copy *PGC::mCherry* marker (Figure 1A) that is driven by the *mex-5* promoter. In contrast to *vbh-1* and *glh-1*, zygotic *mex-5P::*mCherry was expressed at a low and similar level in the PGCs of both starved and fed L1 larvae, as well as in starved *daf-18* mutants (Figure S4B). This suggests that not all germline genes are up-regulated by the loss of *daf-18*.

In summary, our results together with those of Wong et al. (2018) suggest that DAF-18 represses zygotic germline gene activation of a subset of genes in the absence of food.

### Germline identity appears unaffected by *daf-18*

One possible explanation for the elevated zygotic gene expression in PGCs is that loss of *daf-18* may compromise germline identity such that the PGCs express genes zygotically on a schedule similar to the soma. If so, zygotic somatic gene expression might occur in PGCs of *daf-18* mutants. To test this hypothesis, we investigated zygotic expression of *unc-119P::GFP*, a neuronal gene reporter previously shown to be expressed inappropriately in the germ line under conditions where germ cell identity is compromised (Ciosk et al. 2006; Mainpal et al. 2015; Tursun et al. 2011; Updike et al. 2014). We found that zygotic *unc-119P::GFP* is not detectable in PGCs, neither in the starved or fed wild type, nor the starved *daf-18* mutant (Figure S4C). On the other hand, starved wild-type animals show zygotic *unc-119P::GFP* expression in neurons, and the fluorescence intensity of this expression was slightly elevated in *daf-18* mutants. In addition, continuously fed wild type and *daf-18* mutants showed no *unc-119P::GFP* expression in the germ line as adults (data not shown). Finally, in no case did we see the germline-specific *glh-1* or *mex-5* reporters expressed outside of the germ line in *daf-18* mutants. Therefore, within the limits of these assays, we found no evidence for altered germline-soma identity in *daf-18* mutants.

### Transcript levels of a subset of germline genes are elevated in starved daf-18 mutants

We reasoned that if *daf-18* mutant PGCs robustly activate zygotic germline gene expression despite or even prior to starvation, more pervasive effects on germline transcript levels should be detectable. To test this prediction, we used a combined mRNA-sequencing and bioinformatics approach to compare germline and somatic gene expression. We compared transcript levels from whole, starved *daf-18* mutant and wild-type L1 larvae within two hours of hatching, in four independent biological replicates (Figure 5A). We then used available annotation of tissue-specific expression and gene function to analyze the results. Principal Components Analysis and pairwise correlations demonstrated reproducibility of replicates and a relatively large effect of *daf-18* on gene expression (Figure S5-1), with the first principal component clearly separating the two genotypes and explaining over half of the variance. 3,335 genes were significantly differentially expressed (false discovery rate (FDR) < 0.05) (Figure 5B); 1,802 genes were down-regulated and 1,533 up-regulated in *daf-18* mutants relative to the wild type (File S1).

**Figure 5.**
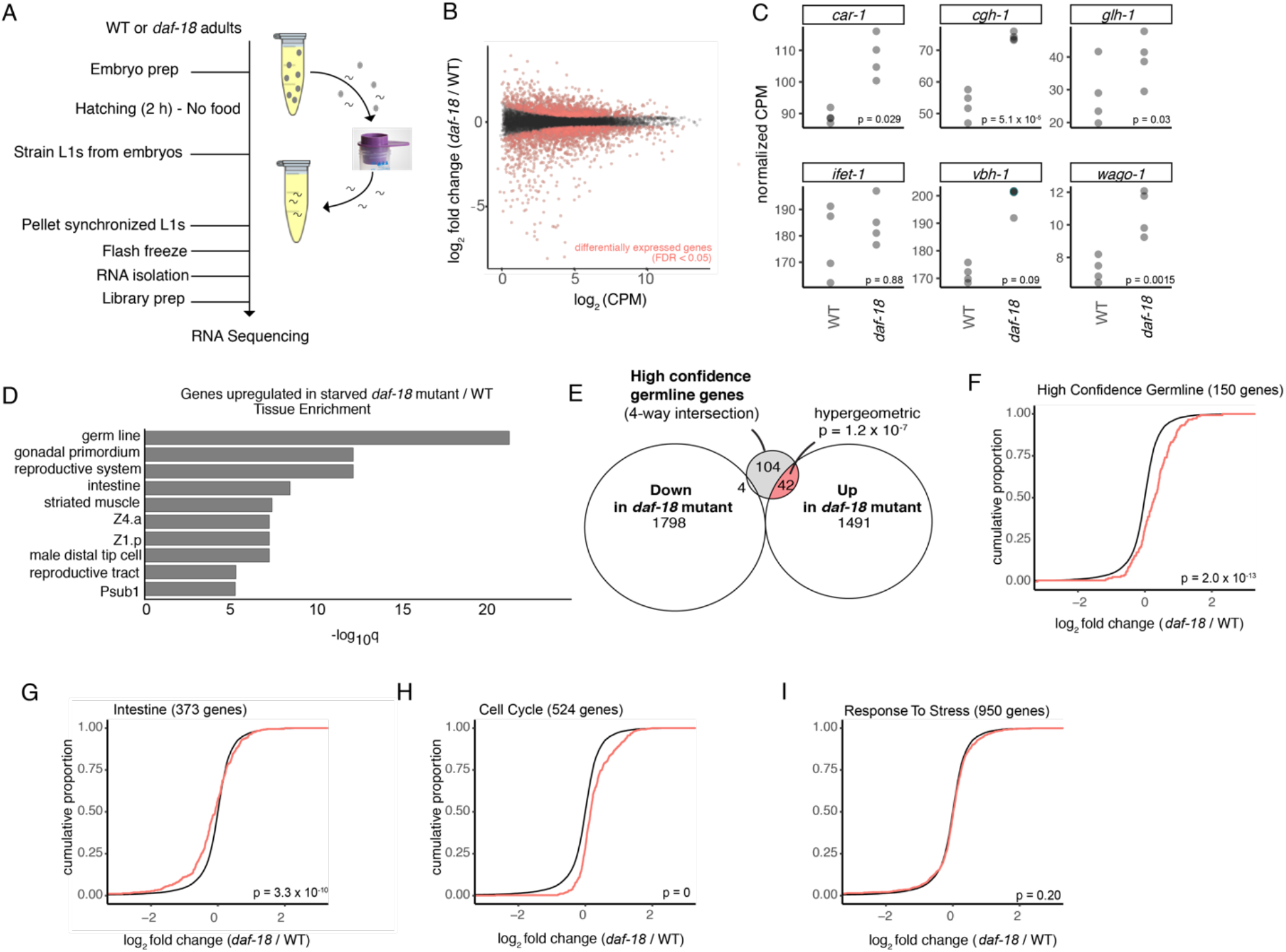
Transcript levels of a subset of germline genes are elevated in starved *daf-18* mutants. **A**. Schematic of RNA Sequencing experiment. Starved L1s were isolated within 2 h of hatching, pelleted, frozen, and mRNA sequencing was performed. **B**. Plot of fold change (in GC1459 *daf-18(ok480)* / GC1171 wild type (WT) versus expression level of 12,592 transcripts detected. Red color dots represent transcripts expressed differentially (below cutoff of FDR 0.05). **C**. Transcript levels of genes previously shown to be induced zygotically by feeding (Wong et al. 2018) or by *daf-18* mutation (this study). Uncorrected p-values are displayed since comparisons were hypothesis-driven. **D**. Tissue enrichment analysis of genes upregulated in *daf-18* mutant/WT using WormBase web tool. Germ line is the most significantly represented tissue in *daf-18* mutant-upregulated genes. **E**. Venn diagram of transcripts significantly upregulated or downregulated in *daf-18* mutants (FDR<0.05) and overlap with the set of “high confidence germline genes,” or transcripts found in all of 4 published germline sets (Cao et al. 2017; Lee et al. 2017; Packer et al. 2019; Spencer et al. 2011). Hypergeometric p-value is displayed for *daf-18* up-regulated genes overlap with high confidence germline genes (while p=0.99999977 for overlap of *daf-18* downregulated genes with high confidence germline genes). **F-I**. Cumulative distribution function (CDF) plots of transcript sets of interest (red line) versus the background set of all 12,592 transcripts detected (black line). Shift to the right indicates overall increased expression of gene set. p-values determined by Kolmogorov-Smirnov (KS) test are shown. Numbers of transcripts from published gene sets detected in our experiment are shown in graph titles (parentheses). **G**. Intestine gene set (Cao et al. 2017), data accessed by GExplore1.4 web tool) shows significant differential expression in *daf-18*, but many genes are downregulated. **H**. and **I**. show significant upregulation (Cell Cycle) and no significant change (Response to Stress) of gene sets of interest.

To focus our comparative analysis on germline genes, we performed several independent analyses. First, we examined the five genes that Wong et al., 2018 had shown by mRNA FISH to be up-regulated upon feeding (*vbh-1, car-1, cgh-1, wago-1, ifet-1*), plus *glh-1* that, like *vbh-1*, had elevated zygotic expression in starved *daf-18* mutants based on live GFP protein fusion reporters (Figure 4). *car-1, cgh-1, wago-1*, and *glh-1* were significantly up-regulated in starved *daf-18* mutants compared to the wild type (Figure 5C). *vbh-1* and *ifet-1* were also elevated, but not to the level of statistical significance. This may be related to their somewhat broader tissue expression in the soma compared to the other four genes (Cao et al. 2017).

Second, we performed an unbiased analysis to determine which tissues were most highly represented by the list of genes upregulated in *daf-18* mutants. Using the Enrichment Analysis web tool provided by WormBase (Angeles-Albores et al. 2016; Angeles-Albores and Sternberg 2018), https://wormbase.org/tools/enrichment/tea/tea.cgi), we found that the tissue that showed the most significant enrichment in the genes for which expression went up in *daf-18* mutants was the germ line (Figure 5D).

Third, to identify a set of high-confidence germline enriched genes, we took advantage of four independent lists of genes previously reported to be enriched or expressed in the germ line (Cao et al. 2017; Lee et al. 2017; Packer et al. 2019; Spencer et al. 2011); see Methods for details). Our experiment detected transcripts of a large percentage of the genes in these sets (Figure S5-2A, File S1 for all genes detected and statistical analyses). We found that these four independent analyses identified 167 germline genes in common, 150 of which were detected in our experiment. We refer to these as the “high confidence germline gene” set. Notably, of the 46 genes in this set that were differentially expressed, all but 4 were up-regulated in the *daf-18* mutant (Figure 5E). In addition, a cumulative distribution function (CDF) plot of all genes in the high confidence germline set also revealed a significant shift toward higher expression for the set of high confidence germline genes in *daf-18* mutants (Figure 5F). Each independent germline gene set (Cao et al. 2017; Lee et al. 2017; Packer et al. 2019; Spencer et al. 2011) also showed a significant shift toward higher expression in *daf-18* mutants (Figure S5-2 B-E).

To determine whether genes expressed in specific somatic tissues were similarly affected we used the GExplore1.4 web tool (Cao et al. 2017; Hutter and Suh 2016) http://genome.sfu.ca/gexplore/gexplore_search_all.html) to select genes expressed in the intestine, ciliated neurons, touch receptor neurons, non-seam hypodermis, seam cells, body wall muscle and pharyngeal gland (all are reported to express *daf-18* except for touch receptor neurons according to GExplore data, and possibly pharyngeal gland cells (Packer et al. 2019)). In each case, RNA sequencing detected the majority of the genes in these lists (File S1). We found significant differential gene expression in *daf-18* versus the wild type for genes expressed in the intestine, ciliated neurons, touch receptor neurons, and non-seam hypodermis. However, as shown by the CDF plots, none of these gene sets showed the same overall shift toward elevated expression as was seen with the high confidence germline gene sets (Figure 5G shows intestine; other tissues are shown in Figure S5-3).

We then asked whether genes associated with relevant GO terms showed differential expression in *daf-18* mutants. We selected GO terms associated with the germ line (Reproductive Process, Germ Cell Development, P granule, Cytoplasmic Stress Granule, Gene Silencing by RNA), as well as terms relevant to *daf-18* and/or PTEN function (Cell Cycle, Response to Stress, DNA Damage, Translation). Finally, we looked at genes annotated with both Translation and Reproductive Process GO terms to assess translation within the germ line. Based on our analysis of CDF plots, we found that genes in each of these GO term categories were up-regulated in starved *daf-18* mutants, with the exception of the “cytoplasmic stress granule” and “response to stress” classes. We were not surprised to see cell cycle genes upregulated (Figure 5H), given that all cells that divide in fed L1 larvae, including germ cells and somatic cells, proliferate inappropriately in starved *daf-18* L1 larvae (Fukuyama et al. 2015; Fukuyama et al. 2006; Zheng et al. 2018). The “cytoplasmic stress granule” class contained only 15 genes, limiting statistical power, so interpretation of this finding is difficult (Figure S5-3 H). However, our failure to detect differential gene expression in the “response to stress” set of 950 genes suggests that the loss of *daf-18* does not appreciably alter the animal’s global stress response at the transcriptional level (Figure 5I). Genes annotated with the other seven GO terms examined were significantly differentially expressed (upregulated overall) (Figure S5-3).

We also performed an unbiased analysis of gene ontology (GO) terms, using WormBase’s Enrichment Analysis Tool. We found that among the genes upregulated in starved *daf-18* mutants, some of the most significantly enriched GO terms were related to metabolism and cell division (Figure S5-3 O).

Taken together, our results from this RNA-Seq experiment, as well as zygotic gene reporter expression experiments, indicate that DAF-18 plays a major role in suppressing expression of germline genes during starvation. This effect appears to be restricted to the germ line, though it is possible that some germline genes are upregulated in the soma.

### Regulation of zygotic germline gene activation by *DAF-18* is partially independent of TOR

Previous results showed that depleting *let-363*/TOR by RNAi significantly suppresses the inappropriate cell division phenotype of *daf-18* mutants (Fukuyama et al. 2012). We wondered whether interfering with TOR by RNAi would similarly suppress the inappropriate zygotic germline gene expression in PGCs of starved *daf-18* mutants. To test this hypothesis, we depleted *let-363*/TOR by RNAi and assayed both PGC division and zygotic expression. We initiated RNAi feeding in the L4 stage of hermaphrodites, crossed the GFP-tagged VBH-1 or GLH-1 fusion from the male, and scored the starved L1 progeny for the number of PGCs and the expression level of the tagged protein. As an additional control for the efficacy of RNAi, we also examined progeny of these matings that had been continuously fed on RNAi or control bacteria in parallel. While the progeny exposed to control bacteria became fertile adults, we observed completely penetrant larval arrest on *let-363* RNAi, mimicking the null mutant phenotype (Long et al. 2002) and thereby confirming good efficacy of the RNAi. We also confirmed that *let-363* RNAi suppresses PGC divisions in starved *daf-18* mutant L1 larvae (Figure 6), as was shown previously (Fukuyama et al. 2012). We also noted that PGC divisions also did not occur in fed wild-type L1 larvae upon *let-363* RNAi within the time frame of our scoring.

**Figure 6.**
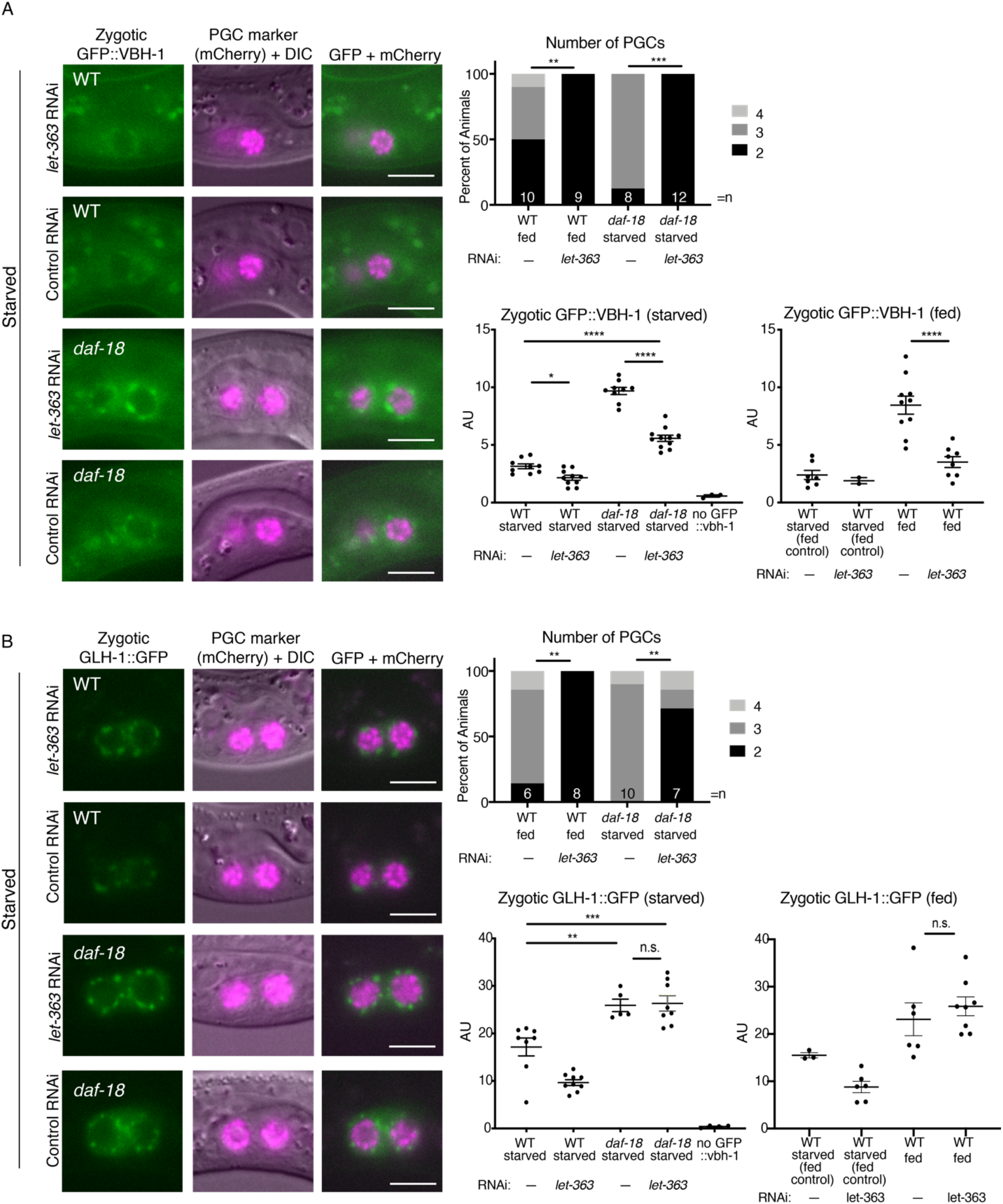
Regulation of zygotic germline gene activation by DAF-18 is partially independent of TOR. **A-B**. Crosses were performed as in Figure 4, but on RNAi bacteria (RNAi by feeding), and L1 progeny analyzed, either starved (images and graph) or fed (graph only). RNAi bacteria carried dsRNA to *let-363*/TOR or Control (empty vector L4440). Images show epifluorescence. Plotted to the right of each set of images are 1) plots of PGC numbers to illustrate efficacy of *let-363*/TOR RNAi in that experiment, and 2) graphs of mean perinuclear GFP fluorescence intensity (AU) values for starved or 3) fed L1 progeny. For PGC number graphs, statistical significance determined by Fisher’s exact test (two-sided). **p<0.01, ***p<0.001. For zygotic GFP images, exposure time is longer for GFP::VBH-1 than GLH-1::GFP. For GFP quantification graphs, each dot represents one animal (either average of 2 PGC values, or one PGC per animal). Non-cross progeny were identified by the lack of somatic GFP (“no GFP::vbh-1”) or lack of any GFP P-granules (“no GLH-1::GFP”; this transgene is much brighter and is clearly visible in starved WT PGCs). Mean +/-SEM. Statistical significance determined by one-way ANOVA (Tukey’s multiple comparisons test).*p<0.05, **p<0.01, ***p<0.001, ****p<0.0001. All scale bars represent 10μm. **A. GFP::VBH-1. “**Starved” L1 progeny were imaged up to 6 hours after hatching (without food). “Starved for fed control” L1 progeny were imaged up to 22 hours after clean embryo preparation (no food), while “fed” L1 progeny were imaged at the same time, but upon 7 hours of feeding (after up to 15 hours post-embryo preparation). Two additional replicates showed similar results (slight suppression of elevated GFP::VBH-1 in *daf-18* mutants, by TOR RNAi). PGC numbers, to assess *let-363*/TOR RNAi efficacy, were counted in L1s starved up to 6 h.**B. GLH-1::GFP**. “Starved” L1 progeny were imaged up to 5 hours after hatching (without food). “Starved for fed control” L1 progeny were imaged up to 22 hours after clean embryo preparation (no food), while “fed” L1 progeny were imaged at the same time, but upon 7 hours of feeding (after 15 hours post-embryo preparation). One additional replicate showed similar results (no suppression of elevated GLH-1::GFP in *daf-18* mutants, by TOR RNAi). PGC numbers, to assess let-363/TOR RNAi efficacy, were counted in L1s starved up to 9 h.

*let-363* RNAi affected zygotic VBH-1 expression in two ways. First, we found that VBH-1 levels were reduced in the PGCs in fed wild-type larvae. Second, the elevation of VBH-1 in *daf-18* mutants relative to the starved wild type was only partially suppressed by *let-363* RNAi (Figure 6A).

We used the same RNAi experimental design to test whether inappropriately elevated GLH-1 zygotic expression in starved *daf-18* mutants is regulated by *let-363/*TOR. We found that, despite strong suppression of PGC divisions in starved *daf-18* mutants, upon *let-363*/TOR RNAi, neither fed WT nor starved *daf-18* mutant animals showed a reduction of zygotic GLH-1 levels (Figure 6B).

Together, our results are consistent with a model (Figure 7) in which DAF-18 impedes zygotic germline gene activation independently of its negative regulation of TOR via opposition of PI3 kinase activity, though both activities ultimately influence PGC division.

**Figure 7.**
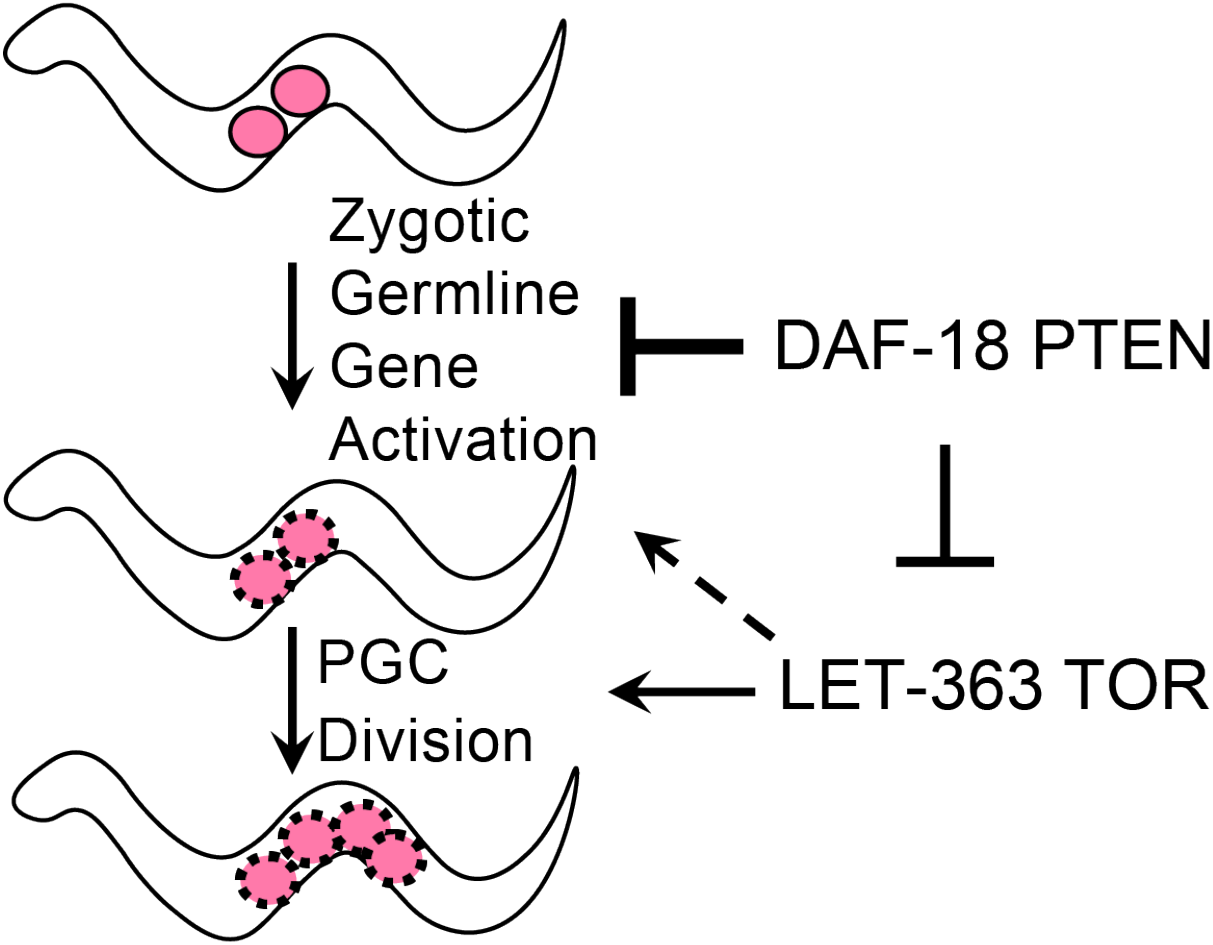
**Model** of DAF-18 repression of zygotic germline gene expression, partially independent of its canonical negative regulation of PI3K-TOR. See text for details.

## Discussion

We investigated several aspects of DAF-18 PTEN function in maintaining PGC quiescence in the absence of food. First, we found that the control of PGC division appears to be remarkably distributed, both between different tissues and also between maternal and zygotic *daf-18* provision. Most striking, we observed nearly complete rescue from a transgene expressing *daf-18(+)* in the germ line and partial suppression from several somatic tissues including the somatic gonad. These findings distinguish the anatomical requirements for *daf-18* in PGC quiescence from its later role in restricting germ cell proliferation during dauer where *daf-18* acts solely in the somatic gonad to regulate germline proliferation (Tenen and Greenwald 2019). Though germ cells arrest in G2 in both cases, this difference underscores differences in regulation of germ cell quiescence in L1 PGCs and later germ cells that have undergone proliferation prior to dauer entry. We also found that either maternal or zygotic *daf-18(+)* appears sufficient to maintain PGC quiescence. While perplexing, one possibility is that this level of redundancy may have evolved as a failsafe to ensure PGC quiescence in the absence of food.

Second, our results point to a role for DAF-18 in activating gene expression in the germ line. Furthermore, they suggest that DAF-18-mediated repression of germline gene activation and PGC cell cycle progression in response to food are not inextricably linked. That is, the role of DAF-18 in activating gene expression in the germ line is partially independent of its role opposing PI3K-TOR signaling.

Third, we found that, within the limits of our assay of *unc-119* expression which is sensitive to germline-to-soma transformation, we found no evidence for *daf-18* activity in regulating the germline-soma identity decision. Nevertheless, the proliferated germ cells in *daf-18* mutants are likely abnormal. For example, in the wild-type, germ cell differentiation occurs upon reduction of GLP-1 Notch signaling in the germ cells. However, the germ cells in starved *glp-1(ts);daf-18* mutant L1 larvae shifted to the restrictive temperature during embryogenesis still divided but did not appear to enter meiosis, unlike those in fed controls (data not shown).

We propose a model in which DAF-18, through as-yet unknown molecular mechanisms, restrains zygotic gene activation in the germ line to interfere with exit from quiescence. Our model rests on the following findings. First, even when worms hatch on food, PGCs of *daf-18* mutant worms initiate divisions earlier than in the wild type, suggesting that they are primed to divide. Second, several germline genes are more highly expressed in PGCs both upon feeding ((Wong et al. 2018); this study) and in starved *daf-18* mutants (this study). Third, we found that several global indicators of active transcription are elevated inappropriately in PGCs of newly hatched *daf-18* mutants, and that some germline genes appear as a group to be upregulated in starved *daf-18* mutant hatchlings relative to the wild type. That said, while only the PGC expression (and not somatic expression) of VBH-1 appeared to be elevated by loss of *daf-18*, our RNA-sequencing results show that transcript levels of several other germline genes were not elevated in *daf-18* relative to the wild type (e.g., *nos-1, nos-2* and *mex-5*; the latter is consistent with our zygotic expression analysis of a *mex-5* driven promoter, Figure S4B). We also note that some somatic transcripts are also elevated in starved *daf-18*, such as *unc-119* (consistent with zygotic transgene expression analysis Figure S4C). Nonetheless, in general, over 90% of differentially regulated genes within the high-confidence set of germline genes are upregulated rather than down-regulated in *daf-18*. Last, elevated levels of zygotic germline gene expression in PGCs of starved *daf-18* mutants occurs even when *let-363* TOR is depleted.

Our model raises many questions for future studies. For example, do the cell cycle and transcriptional regulatory activities of *daf-18* on L1 germ cell quiescence act from the same tissues? Do they result from different subcellular localization and/or biochemical activities of DAF-18 PTEN? For example, in neuroblast divisions, DAF-18 acts via PI3K but upstream of MAPK (Zheng et al. 2018) whereas in oocyte maturation, DAF-18 acts downstream of the VAB-1 Ephrin receptor and upstream of MAPK, independent of AKT and lipid-phosphatase activity (Brisbin et al. 2009). It is possible that the different biochemical activities of DAF-18 correlate with their anatomical requirements. For example, the lipid phosphatase role of DAF-18 PTEN could correlate with post-transcriptional cell cycle control via regulation of PI3K signaling and TOR dependence while a germ cell autonomous function of DAF-18 PTEN (perhaps as a protein phosphatase) could responsible for the transcriptional activation in the germ line. Likewise, the maternal or zygotic requirements may be functionally separable. Additional studies are required to address these questions.

How does DAF-18 interfere with zygotic germline gene activation in PGCs? And what releases this block? The trigger(s) for zygotic germline gene activation are mysterious. In worms, flies, fish, frogs and mammals, zygotic germline gene activation occurs after zygotic somatic genome activation, and, similar to zygotic activation in the soma, it likely requires both degradation of maternal mRNAs and activation of a zygotic program of transcription. Degradation of maternal mRNAs is an important step in securing germ line identity in *C. elegans* (Lee et al. 2017).

While future work is required, several general possibilities can be imagined, none of which are mutually exclusive. Regardless of whether DAF-18 acts biochemically as a lipid phosphatase or protein phosphatase, or otherwise, a downstream effect of DAF-18 could alter one or more processes that promote zygotic germline gene expression. One possibility is that DAF-18 changes the activity state or localization of one or more transcription factors. We noticed, for example, that according to the WormExp (Yang et al. 2016), 37 of the 46 genes that were differentially expressed in *daf-18* mutants are also down in *ceh-23* mutants (Chang et al. 2017). Another hypothesis that could account for the large number of activated germline genes is that DAF-18 may interact with chromatin, DNA, or components of transcription complexes, including RNA PolII, to alter transcription at specific loci, as proposed for nuclear PTEN in mammalian cell culture (Steinbach et al. 2019). DAF-18 may also have a more global effect on chromatin conformation, either via histone modifications or regulation of the degree of physical compaction of chromatin. This hypothesis is intriguing since the timing of chromatin decondensation correlates with zygotic germline gene activation in *C. elegans* PGCs in response to feeding (Butuci et al. 2015; Wong et al. 2018). There is also precedent in mammalian cell lines for PTEN involvement in chromatin condensation via interaction with linker histone H1 and stabilization of HP1alpha, both independent of its lipid phosphatase activity (Chen et al. 2014; Gong et al. 2015). Condensed chromatin may be an evolutionarily conserved mechanism for securing the quiescent state, as it is also implicated in yeast and human cell lines (Swygert et al. 2019). Small RNAs and nuclear RNAi factors were recently shown to induce chromatin compaction in later stage germ cells (Fields and Kennedy 2019; Weiser et al. 2017)). We speculate that the same mechanisms used by DAF-18 to restrict zygotic germline gene expression in *C. elegans* could conceivably contribute to the competence of cells arrested in G2 to escape quiescence in the absence of PTEN in cancer.

## Methods

### Worm Handling and Strains

*C. elegans* strains were maintained on nematode growth medium (NGM) plates at 20°C and fed a diet of OP50 *E. coli* bacteria, unless noted otherwise (Stiernagle 2006). Experiments were carried out at room temperature (∼22°C).

#### Strains used in this study

Strains generated for this study are indicated below. All strains used and details on strain construction information are listed in Table S1. **GC1459** *naSi2 II; unc-119(ed3) III ?; daf-18(ok480) IV*, **GC1527** *glh-1(sam24[glh-1::gfp::3xFLAG]) I; daf-18(ok480) IV*, **GC1537** *vbh-1(na110[GFP::vbh-1]) I; naSi2 II*, **GC1576** *vbh-1(na110[GFP::vbh-1]) I; naSi2 II; daf-18(ok480) IV*, **GC1456** *naSi2 II; unc-119(ed3) III ?; daf-18(ok480) IV; naEx261 (daf-18P::daf-18::daf-18 3’UTR::SL2::GFP::H2B::unc-54 3’UTR)*, **GC1460** *naSi2 II; unc-119(ed3) III ?; daf-18(ok480) IV; naEx264 (rab-3P::daf-18::daf-18 3’UTR::SL2::GFP::H2B::unc-54 3’UTR)*, **GC1464** *naSi2 II; unc-119(ed3) III ?; daf-18(ok480) IV; naEx268 (nlp-40P::daf-18::daf-18 3’UTR::SL2::GFP::H2B::unc-54 3’UTR)*, **GC1465** *naSi2 II; unc-119(ed3) III ?; daf-18(ok480) IV; naEx269 (dpy-7P::daf-18::daf-18 3’UTR::SL2::GFP::H2B::unc-54 3’UTR)*, **GC1485** *naSi2 II; unc-119(ed3) III ?; daf-18(ok480) IV; naEx275 (elt-2P::daf-18::daf-18 3’UTR::SL2::GFP::H2B::unc-54 3’UTR)*, **GC1484** *naSi2 II; unc-119(ed3) III ?; daf-18(ok480) IV; naEx274 (ehn-3P::daf-18::daf-18 3’UTR::SL2::GFP::H2B::unc-54 3’UTR)*, **GC1479** *naSi22 (mex-5P::daf-18::nos-2 3’UTR::SL2::GFPo::PH::tbb-2 3’UTR + unc-119(+)) II; unc-119(ed3) III ?; daf-18(ok480) IV*, **GC1483** *glh-1(sam24[glh-1::gfp::3xFLAG]) I; naSi22 (mex-5P::daf-18::nos-2 3’UTR::SL2::GFPo::PH::tbb-2 3’UTR) II; unc-119(ed3) III ?; daf-18(ok480) IV*

Transgenic strains were generated by three different methods, as indicated below (see Table S1):

1. Microinjection (generating extrachromosomal arrays) (Evans 2006) directly into GC1459 *naSi2 II; daf-18(ok480) IV*. All arrays used pJB134 (*flp-17*P::dsred) as a co-injection marker (Brandt and Ringstad 2015)
2. Cas9-triggered long-range HDR using plasmid repair template, and plasmid Cas9 and single guide RNA (Dickinson et al. 2013). Although the *Phsp::peel-1* was included in the injection mix we found that the heat shock step was not necessary (Used to generate *germline::daf-18 (naSi22)*).
3. CRISPR-based “hybrid” GFP donor method (Dokshin et al. 2018), with minor modifications from J. Nance that used *dpy-10* co-CRISPR instead of *rol-6(su1006d)*. Injection was done into *GC1171 naSi2 II* to generate *GFP::vbh-1 (na110)*. crRNA targeting sequence for *vbh-1* and *dpy-10* (co-CRISPR) were generated by IDT. https://www.idtdna.com/site/order/designtool/index/CRISPR_CUSTOM. Cas9 and tracrRNA were purchased from IDT.

### Starved L1 preparation / L1 synchronization

#### Loosely-synchronized L1s (From small scale embryo preparation)

For some experiments (Images in Figure 1A, tissue-specific *daf-18* “rescue” experiments in Figure 2, a-amanitin experiment in Figure 3E, and immunofluorescence in Figure S3B) a loose synchronization was sufficient. For this protocol, each strain was grown on 1-3 6 cm NGM plates seeded with OP50. Gravid hermaphrodites and embryos were washed with ∼1 mL M9 buffer from plates into a 1.8 mL Eppendorf tube. A pure preparation of clean (bacteria-free) embryos was obtained by sodium hypochlorite treatment (1 mL, freshly-prepared, final concentration: 10% sodium hypochlorite solution [Sigma #425044, available chlorine 10-15%], 10% 5N NaOH, 80% M9 buffer). Reactions were monitored under a dissecting microscope and proceeded until no whole adult carcasses remained (∼2-5 min). Samples were then washed 5x with ∼1 mL M9 buffer (with pelleting at ∼1690+ g), yielding purely embryos. Embryos hatched overnight in ∼1 mL M9 buffer + 0.08% ethanol in 1.8 mL tubes, with rocking (rocking was omitted for a-amanitin experiment). Therefore, L1 larvae were loosely synchronized with respect to hatching. L1 larvae were assessed at ∼24 h or ∼72 h from the embryo preparation as noted in figure legends, or in graph titles for Figures 2 and 3E, as “1 Day Starved” or “3 Days Starved”, respectively.

#### Tightly-synchronized L1s (From large scale embryo preparation)

For experiments requiring tight synchronization of large batches of L1 larvae with respect to hatching (experiments: timing of PGC divisions in Fig 1B-C, immunofluorescence in Figure 3, and RNA Sequencing in Figure 5), animals were grown on a larger scale for embryo preparation. Each strain was grown on 4-6 high peptone NGM plates (10 cm) seeded with OP50 (starting with 40-60 adult hermaphrodites per plate and growing for 5 days at 20°C). Gravid hermaphrodites and all embryos were washed with 10 mL M9 buffer into 2-3 conical tubes (15mL). Pure preparations of clean (bacteria-free) embryos were obtained by sodium hypochlorite treatment (10 mL, freshly-prepared, final concentration: 10% sodium hypochlorite solution [Sigma #425044, available chlorine 10-15%], 10% 5N NaOH, 80% M9 buffer). Reactions were monitored under a dissecting microscope and proceeded until no whole adult carcasses remained (∼2-5 min). Samples were then washed 5x with ∼10mL M9 buffer (with pelleting at ∼1690 g), yielding purely embryos. Embryos hatched for 2h in 10 mL M9 buffer (Immunofluorescence experiments, Figure 3) or M9 + 0.08% ethanol (100%) (timing of PGC divisions in Fig 1B-C and RNA Sequencing in Figure 5), in 2-3 conical tubes (15 mL). Resulting L1s were separated from embryos using 10 μm mesh Pluristrainers (Pluriselect # 43-50010-03). L1s were strained into 50 mL conical tubes (and embryos caught in the filter), washed 5x with 5mL M9, with swirling and pipetting up and down to maximize L1 yield and prevent clogging of filter with embryos. Resulting L1s larvae were tightly synchronized, having hatched within the previous 2 hours. To proceed with immunostaining or RNA Sequencing, larvae were pelleted by centrifugation for 2 min at 900-1300g (yields ∼5 μL or less of compacted L1s, or estimated 20,000-40,000 worms). For RNA Sequencing experiment, pelleted L1 larvae were flash frozen in liquid nitrogen and stored at −80°C until shipping all samples on dry ice for analysis.

#### Tightly-synchronized L1s (from very small scale embryo preparation)

For experiments requiring selection of individual hermaphrodite mothers (Maternal/Zygotic *daf-18* requirement experiments (Figure 1B-C) and all zygotic gene expression experiments (Figures 4, 4S, 6, 6S), gravid hermaphrodite P0s were selected and then bleached on unseeded NGM plates. Up to 15 hermaphrodites were placed in the center of the plate and treated with ∼15μL of 1:1 sodium hypochlorite solution (Sigma, 10-15% available chlorine) and 5N NaOH. Each reaction was monitored under a dissecting microscope, and free embryos were moved out of the bleach drop with a sterile worm pick, while adult carcasses and fragments remained in the bleach. Once all carcass fragments were dissolved and bleach was absorbed into the plate, 15-30μL of M9 buffer was placed on top of embryos as a “wash.” Most embryos had not hatched ∼12 hours later, so hatching of remaining embryos could be monitored hourly thereafter (zygotic gene expression experiments) or after several days (maternal/Zygotic *daf-18* requirement experiments).

### Timing of hatching from 2-4 cell stage

Embryos were isolated by dissection of gravid hermaphrodites in M9 in glass wells. Embryos at the 2-4 cell stage were selected under the dissecting microscope with a mouth pipette and transferred to unseeded NGM plates. Embryos were monitored hourly for the number of hatched L larvae and embryos in 4 replicates (2 biological, 2 technical).

### Timing of PGC divisions

Tightly-synchronized L1s were isolated using the large-scale embryo preparation (hatched within a 2h time frame, described above). The timing in Figures 1B and 1C are labelled as “Hours post-collection,” meaning that animals examined at time point “0” hatched within the last 2 hours. L1 animals were either left in liquid culture (Figure 1B, “starved”) or plated on NGM plates seeded with OP50 bacteria (Figure 1C, “fed”) for the designated amounts of time (1-4 h).

### PGC counts

L1 larvae were mounted onto 2% agarose pads on slides, paralyzed with 1mM levamisole (Sigma), and imaged on a Zeiss Z1 Axioimager (imaged with 63x objective) within 1 hour of mounting. PGC number was determined by counting *naSi2+ (PGC::mCherry)* fluorescent nuclei, or for *germline::daf-18* “rescue” experiments, PGCs with GLH-1::GFP perinuclear GFP or membrane *GFP* (*germline::daf-18::SL2::GFP::PH*). Dead L1 larvae were always omitted; these were identified by a distinctive auto-fluorescent glow and/or a highly vacuolated appearance.

### Immunofluorescence

Strains bearing the PGC marker GLH-1::GFP, fluorescence from which generally survived fixation, were used for all immunofluorescence experiments except for Figure S3A (starved L1 larvae, stained with anti-PGL-1 antibody OIC1D4).

#### Partial dissociation/permeabilization

L1 larvae were first prepared by “Tight Synchronization/Large-scale preparation” described above, which yields a small L1 pellet (∼5μL or less of compacted L1 larvae) (for staining experiments in Figure 3). Loosely-synchronized L1 larvae (from overnight hatching of clean, washed embryos prepared by small or large-scale embryo preparation, described above), can also be permeabilized this way (Figure S3B). Note that the compacted L1 pellet should be 40μL or less. Larger pellets appear to permeabilize faster, so adjustments to treatment time and strength of mechanical disruption may be required. Similar adjustments may be required for staining of head or tail cells, which are occasionally lost with this permeabilization method. L1s were chemically and mechanically permeabilized by following an existing L1 cell isolation protocol, which details reagents and recipes for buffers required (Zhang and Kuhn 2013) http://www.wormbook.org/chapters/www_cellculture/cellculture.html)), with minor modifications described below.

Briefly, the permeabilization with modifications was performed as follows (a more detailed protocol is available upon request): The L1 pellet was washed with 1mL ddH_2_O in a low retention Eppendorf tube and re-pelleted at 16,000g for 2 min after which as much supernatant as possible was removed with careful pipetting. 200μL of freshly-thawed SDS-DTT solution (20 mM HEPES pH 8.0, 0.25% SDS, 200 mM DTT, 3% sucrose) was added and larvae were incubated for exactly 2 min at room temperature, after which 800μL of egg buffer was added and mixed gently. L1 larvae were then pelleted at 16,000g for 1 min, and washed 5x with egg buffer and gentle mixing; supernatant was again carefully removed. 100μL of freshly-thawed pronase E (15mg/mL, Sigma P8811 in egg buffer) was added and larvae incubated for exactly 7 min at room temperature. During this incubation, a 25 1/2 gauge needle, attached to a syringe pump, was used to suck up-and-down the entire worm suspension a few times each minute. Finally, 900μL L15-FBS was added to stop the reaction (egg buffer may also work). L1 larvae and fragments were then pelleted for 3 min at 1300g, and washed 2x with L15-FBS and gentle mixing. After the last wash, the supernatant was removed, leaving ∼100μL of L1 larvae and fragments in buffer.

#### Fixation

For H3K4 antibodies (and anti-PGL-1, OIC1D4, used to co-stain in one experiment), methanol/acetone fixation was conducted in Eppendorf tubes. Fresh aliquots of methanol and acetone were pre-chilled at −20°C. L1 pellet was gently mixed in residual ∼100 μL buffer, 1 mL cold methanol was added, and tubes were incubated for 9 minutes at room temperature or - 20°C. Samples were centrifuged for 1 min at 16,000 g and supernatant removed, leaving ∼100 μL or less. L1s were resuspended in residual methanol by pipetting and/or brief vortexing (getting as close to single worm suspension as possible to reduce clumping when acetone is added). 1 mL of cold acetone was added, and tubes were incubated for 4 minutes at room temperature or −20°C. Samples were centrifuged for 1 min at 16,000 g and supernatant removed, leaving ∼100 μL or less, then washed 3x with PBS-Tw (PBS with 0.1% Tween). Each wash was 5-10 min with rocking, or agitation by hand several times during the incubation.

For H5 staining, methanol/formaldehyde fixation was used. The protocol above was followed, with the following changes: −20°C methanol step was 2 min, followed by formaldehyde fix (1x PBS, 0.08M HEPES (pH 6.9), 1.6 mM MgSO4, 0.8mM EGTA, 3.7% formaldehyde) (Seydoux and Dunn 1997) for 5 min at room temperature. Wash buffer was PBS-Tr (PBS with 0.1% Triton-X).

#### Antibody staining

For H3K4me antibodies (and anti-pgl-1 OIC1D4), blocking was with 100 μL (or more) PBS-Tw + 0.1% BSA for 30 min. For H5 staining, blocking was with 100 μL (or more) of 9 parts PBS-Tr + 0.1% BSA to 1 part normal goat serum (NGS) for 30 minutes.

1° antibodies, all diluted in respective blocking buffers and incubated overnight at 4°C, were: CMA303 (mouse anti-H3K4me2, Millipore Sigma, Cat # 05-1338-S, diluted 1:20) (Bowman et al. 2013; Furuhashi et al. 2010), ab32356 (rabbit anti-H3K4me2, Abcam, diluted 1:250) (Wang et al. 2011), ab8580 (rabbit anti-H3K4me3, Abcam, diluted 1:500) (Xiao et al. 2011), H5 (mouse anti-P-Ser2 RNA Pol II, Biolegend Cat. # 920201, diluted 1:50), and OIC1D4 (mouse anti-PGL-1, Developmental Studies Hybridoma Bank, gift of J. Nance, diluted 1:2). After 1° antibody incubation, samples were washed for 10 min with rocking, 3-5x with PBS-Tw or PBS-Tr.

2° antibodies used, diluted in respective blocking buffers, were: Goat anti-mouse IgG antiserum, Cy3 conjugated (Jackson Labs, Cat. # 115-165-164, diluted 1:200), goat anti-rabbit IgG CY3-conjugated (from Jackson Labs in 2012, Cat. # 111-166-003, diluted 1:200), goat anti-mouse IgM-Alexa 594 (Invitrogen, Cat. # A-21044, diluted 1:200).

After 2° antibody incubation, samples were washed for 10 min with rocking, 3-5x with PBS-Tw or PBS-Tr with DAPI (25 μg/mL) in the second wash.

Stained L1s were mounted on 2% agarose pads with ProLong Glass antifade mounting media, covered with coverslips, and analyzed immediately, or allowed to cure overnight for imaging the next day.

### alpha-amanitin treatment

L1 larvae were loosely synchronized (using small scale embryo prep) and treated with 10 μg/mL alpha-amanitin (Sigma A2263), as in Butuci et al. (2015). These experiments did not utilize rocking during L1 starvation. Animals that appeared dead by bright auto-fluorescence throughout the body, or highly vacuolated appearance, were omitted.

### Maternal and zygotic *daf-18* requirements

#### Maternal-Zygotic+

*cdc-42::GFP* was crossed in from the male (P0), as a marker to identify cross-versus self-progeny, along with a copy of the *daf-18(+)* gene (genomic, wild type). Hermaphrodite mothers of the cross (P0) were *naSi2 (PGC::mCherry); daf-18(ok480)*. Animals mated for ∼24 hours on NGM seeded with OP50 “cross plates” (small bacterial lawn). Mated hermaphrodites were treated with sodium hypochlorite on unseeded NGM plates (Very Small Scale L1 synchronization, described above) to isolate embryos free of bacteria. Progeny (starved L1 larvae) were scored 3 days later and for numbers of PGCs and presence or absence of *cdc-42::GFP* (inferred *daf-18* (+) positive/negative).

#### Maternal+ Zygotic-

*daf-18(0)* gravid hermaphrodite mothers carrying a *daf-18(+)* genomic rescuing array (*naSi2; daf-18(ok480); naEx261 (daf-18P::daf-18::SL2::GFP)*+) were selected by fluorescence (both *daf-18 GFP* positive and co-injection/transformation marker, *flp-17P::dsRed* positive). These worms were treated with sodium hypochlorite on an unseeded NGM agar plate (Very Small Scale L1 synchronization, described above) to isolate embryos free of bacteria. Resulting L1 worms (starved) were assessed 3 days later, scoring for the number of PGCs and presence or absence of the rescuing transgene (only animals lacking both the transformation marker and *daf-18 GFP* were counted as negative).

### Zygotic gene expression

Males carrying homozygous fluorescently-tagged genes (*na110 [GFP::vbh-1], sam24 [glh-1::GFP], naSi2 [mex-5P::mCherry]*, and *otIs45 [unc-119P::GFP]*) were generated by heat shock, and populations were expanded and maintained by crossing to hermaphrodites of the same strain. 2-4 crosses were performed per strain, in both the *daf-18(+)* and *daf-18* mutant backgrounds, with 25-30 males and 8-10 hermaphrodites per cross. Hermaphrodites carried an alternate-color PGC marker (e.g. *glh-1::GFP* males crossed to *mex-5::mCherry* hermaphrodites). P0 animals mated for 24 hours on NGM plates with small lawns of OP50, and then mated hermaphrodites were treated with sodium hypochlorite on unseeded NGM plates to isolate clean embryos (Very small scale embryo preparation). 12-16 h later, unhatched embryos (mostly 3-fold stage) were moved to fresh unseeded NGM plates by mouth pipetting with M9 buffer. Starved L1 animals were then analyzed for fluorescent zygotic gene expression within as short as 1 h from hatching (and up to 5 h). For fed WT L1 analyses, starved L1 progeny (from crosses that had hatched overnight on the unseeded NGM plates;12-19h from sodium hypochlorite treatment of P0 animals), were moved to OP50 bacterial lawns on NGM plates by mouth-pipetting with M9 buffer. After 5 hours of feeding, these animals were analyzed for zygotic gene expression (fluorescence), along with “starved (fed control)” L1 larvae that remained on unseeded NGM plates (starved up to 17-24h).

For zygotic gene expression experiments utilizing RNAi, crosses were performed on standard RNAi plates (IPTG/Amp) seeded with small lawns of RNAi bacteria (HT115 carrying plasmids for dsRNA production) (Kamath et al. 2003; Timmons et al. 2001). Hermaphrodites fed on RNAi bacteria starting from the L4 stage until 24 hours later (adulthood), and were then treated with sodium hypochlorite on NGM plates to isolate clean embryos (Very small scale embryo preparation). The negative control was the empty vector (L4440). *let-363* RNAi efficacy was confirmed by suppression of *daf-18* PGC divisions in starved L1 progeny, and 100% larval (L3) arrest in progeny from crosses that had continued feeding on RNAi plates (Figure 6). RNAi efficacy for additional experiments (2 replicates for *GFP::vbh-1*, one replicate for *glh-1::GFP*) was confirmed only by 100% larval arrest in RNAi-fed progeny.

### Microscopy and Image Analysis

Imaging was performed with a Zeiss Z1 Axio Imager equipped with an AxioCamMRm digital camera, ApoTome for optical sectioning, a plan apo 63X/1.4 oil DIC objective, and Zeiss filters for DAPI, CY3, and eGFP.

#### For tissue-restricted daf-18 GFP expression

Tissue expression of *daf-18 GFP* transgenes in L1 larvae starved for 1-3 days was documented by taking epifluorescence image stacks of animals mounted on 2% agarose pads, paralyzed with 1mM levamisole. Images are shown as Z-projections (Figure S2).

#### For immunofluorescence experiments

Fluorescence image stacks were taken using Apotome for sectioning (63x objective, ∼0.3μm slices, or optimal slice for fluorophore/filter). Exposure times were determined by wild-type stained animals and held constant for all samples and replicate experiments. Mean fluorescence was quantified with Fiji/ImageJ by drawing an ellipse around each PGC (identified by PGC marker GLH-1::GFP or *anti-pgl-1* co-staining), in the image slice where it (based on surrounding P granules (GLH-1::GFP)) was most in focus. The nearest somatic cell/somatic gonad precursor (SGP) mean fluorescence, and background within the worm but outside of nuclei, was measured in the same image. PGC and SGP fluorescence values were corrected for background fluorescence, and PGC/SGP ratio was calculated.

#### For zygotic gene expression experiments

L1s were mounted on 4% agarose pads containing 10-100mM levamisole in M9. Higher levamisole concentrations were used for zygotic transgene expression requiring longer exposure times (*naSi2 [PGC::mCherry]*). GFP, mCherry, and DIC channels were collected for each animal, within 1h of mounting. Exposure times were held constant for each individual experiment but varied based on zygotic transgene. When possible, both PGCs were imaged for each animal and values were averaged, resulting in a single fluorescence value for each animal. If one PGC was obscured (e.g., by intestine/auto-fluorescent gut granules or other cells), it was omitted, and only one PGC was imaged (Figures 4 and 6).

To analyze perinuclear GFP::VBH-1 or GLH-1::GFP expression, a script for Fiji/ImageJ was written by Michael Cammer (NYU Langone Microscopy Laboratory). This allowed identification of PGC nuclei by manual thresholding of each image in the red channel (PGC::mCherry), and automatic measurement of mean GFP channel fluorescence intensity of a 1μm-wide ring surrounding the nucleus.

For zygotic *naSi2* (*PGC::mCherry*) expression, regions of interest were hand drawn around PGC nuclei and mean fluorescence measured.

For *unc-119P::GFP* head fluorescence measurements, ellipsoid regions of interest were drawn over a single neuron in the head (dorsal, just posterior to the nerve ring, likely a chemosensory neuron) and a single tail neuron near the rectum, and mean fluorescence was measured. The lack of any visible *unc-119P::GFP* in PGCs (even upon long exposure times) was noted in all animals that were imaged and measured for head fluorescence.

For all zygotic transgene expression analyses, background fluorescence was measured for each animal in a region within the worm but outside of fluorescent tissues, and was subtracted from zygotic fluorescence-of-interest mean values.

### RNA Sequencing Experiment

#### GC1171 (*naSi2, PGC::mCherry*) versus GC1459 (*naSi2, PGC::mCherry; daf-18 (ok480)*)

##### RNA isolation and RNA-seq Library Preparation

RNA was isolated using TRIzol Reagent (Thermo-Fisher Scientific) following the manufacturer’s protocol with exceptions noted below. The procedure was scaled down in a linear fashion, using only 100 uL Trizol. 5 ug linear polyacrylamide (Sigma-Aldrich) was included as a neutral carrier for RNA precipitation. RNA was eluted in nuclease-free water and incubated at 55°C for approximately four minutes to resuspend the pellet. 140 ng of total RNA was used for each library preparation. NEBNext Ultra II RNA Library Prep kit (New England Biolabs) was used to perform poly-A selection and prepare libraries for sequencing. The final libraries were enriched with 11 cycles of PCR. Libraries were then sequenced using Illumina NovaSeq 6000 to obtain 50 bp paired-end reads.

##### Differential Expression Analysis of RNA-seq data

Version WS273 of the *C. elegans* genome was used to map sequencing reads (file name c_elegans.PRJNA13758.WS273.genomic.fa downloaded from WormBase). Bowtie version 1.2.2 (Langmead et al. 2009) was used to map 50 bp paired-end reads with the following settings: bowtie -I 0 -X 500 -k 1 -m 2 -S -p 2. Mapping efficiency and number of mapped reads can be found as part of File S1. HTSeq version 0.11.2 (Anders et al. 2015) was used to count reads mapping to the WS273 canonical geneset (file name c_elegans.PRJNA13758.WS273.canonical_geneset.gtf downloaded from WormBase). edgeR version 3.24.3 (Robinson et al. 2010) was used for differential expression analysis. Count data was restricted to include only protein-coding genes (20,127 genes). Prior to differential expression analysis, this list was restricted to include only genes with counts-per-million (CPM) greater than one in at least four libraries (12,592 genes). Principal component analysis (PCA) was performed on log_2_ mean normalized CPM values for the genes included in differential expression analysis. PCA separates libraries based on strain (Figure S5-1 A). The Pearson correlation coefficient for each pair of libraries was calculated using log_2_ normalized CPM values for genes included in differential expression analysis. All replicate libraries have a Pearson correlation coefficient of 0.98 – 0.99 (Figure S5-1 B). In edgeR, the “calcNormFactors” function was used to normalize for RNA composition and the tagwise dispersion estimate was used for differential expression analysis. The exact test was implemented in edgeR for pairwise comparison of four biological replicates of GC1171 to four biological replicates of GC1459. 3,335 genes are differentially expressed at an FDR cutoff of 0.05. All edgeR differential expression output, including gene name, log_2_ fold change, expression level, and FDR, can be found as part of File S1. Raw and processed RNA-seq data are available through GEO NCBI at accession number GSE151299.

##### Gene Group Analysis for RNA-seq

We collected several independent lists of genes previously reported to be enriched or expressed in the germ line:

1. 1346 genes enriched in sorted, labeled embryonic primordial germ cells, Z2 and Z3, over somatic blastomere genes (Supplementary File 8 from Lee et al., 2017)
2. 979 genes enriched in sorted, labeled embryonic primordial germ cells, Z2 and Z3, over all embryonic cells (Supplemental File Z2/Z3 #2 from Spencer et al., 2011)
3. 306 genes 5x enriched in the L2 germline (Cao et al. 2017), data accessed by GExplore1.4 web tool for tissue expression (Hutter and Suh 2016)
4. 4980 genes expressed in embryonic primordial germ cells, Z2/Z3, for which all three “pseudotime” bins are above 0 TPM.

To determine whether gene groups of interest (Cao et al. 2017; Lee et al. 2017; Packer et al. 2019; Spencer et al. 2011) tend to be up-or down-regulated in the *daf-18* mutant compared to wild-type, the cumulative distribution of log_2_ fold changes for each group of genes in the GC1459 / GC1171 comparison was plotted along with the cumulative distribution of log_2_ fold changes of all 12,592 detected genes in the GC1459 / GC1171 comparison. Differences between the two cumulative distributions were assessed using the Kolmogorov-Smirnov test. In a complementary approach, the overlap between each gene group of interest and differentially expressed genes that were up-or down-regulated in the GC1459 / GC1171 comparison (FDR < 0.05) was assessed, and the hypergeometric test was used to determine significant overlap. Gene groups used in this analysis can be found in File S1. Finally, tissue enrichments and GO terms were assessed among genes up-and down-regulated in GC1459 / GC1171 using the WormBase Gene Set Enrichment Analysis tool (Angeles-Albores et al. 2016; Angeles-Albores and Sternberg 2018).

## Acknowledgements

Funding for this work was provided by NIH R01GM061706, R01GM130152, R35GM134876 and NYSTEM IIRP C029561 to EJAH, R01 GM117408 to LRB, and ACS 132083-PF-18-029-01-DDC postdoctoral award to ALF. We thank Michael Cammer for assistance with image analysis; the NYU Langone Microscopy Laboratory is partially supported by the Cancer Center Support Grant P30CA016087. We thank members of the Hubbard and Nance labs for discussions, protocols and reagents, Dustin Updike for strains, and Niels Ringstad for plasmids. We thank WormBase. Some strains were provided by the CGC, which is funded by NIH Office of Research Infrastructure Programs (P40 OD010440).

