## Supplemental Figures and Legends for "DAF-18/PTEN inhibits germline zygotic gene activation during primordial germ cell quiescence"

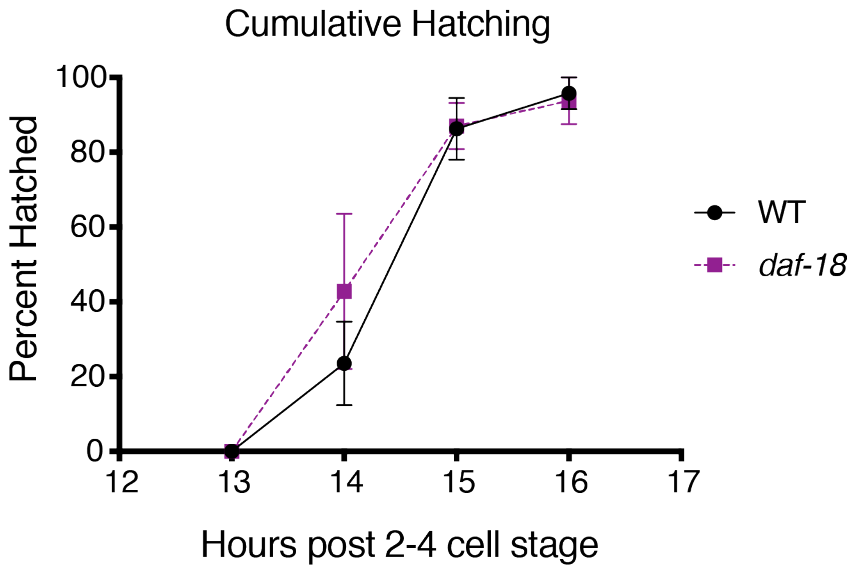


**Figure S1.** ***daf-18*** **mutant embryos hatch at the same time as wild type.**

The number of L1 larvae hatched from embryos isolated at the 2-4 cell stage was monitored hourly. 4 replicate experiments (2 biological, 2 technical), containing a total of 40 WT (*PGC::mCherry*, *naSi2*) animals, and 49 *daf-18* mutant animals (*naSi2; daf-18*).

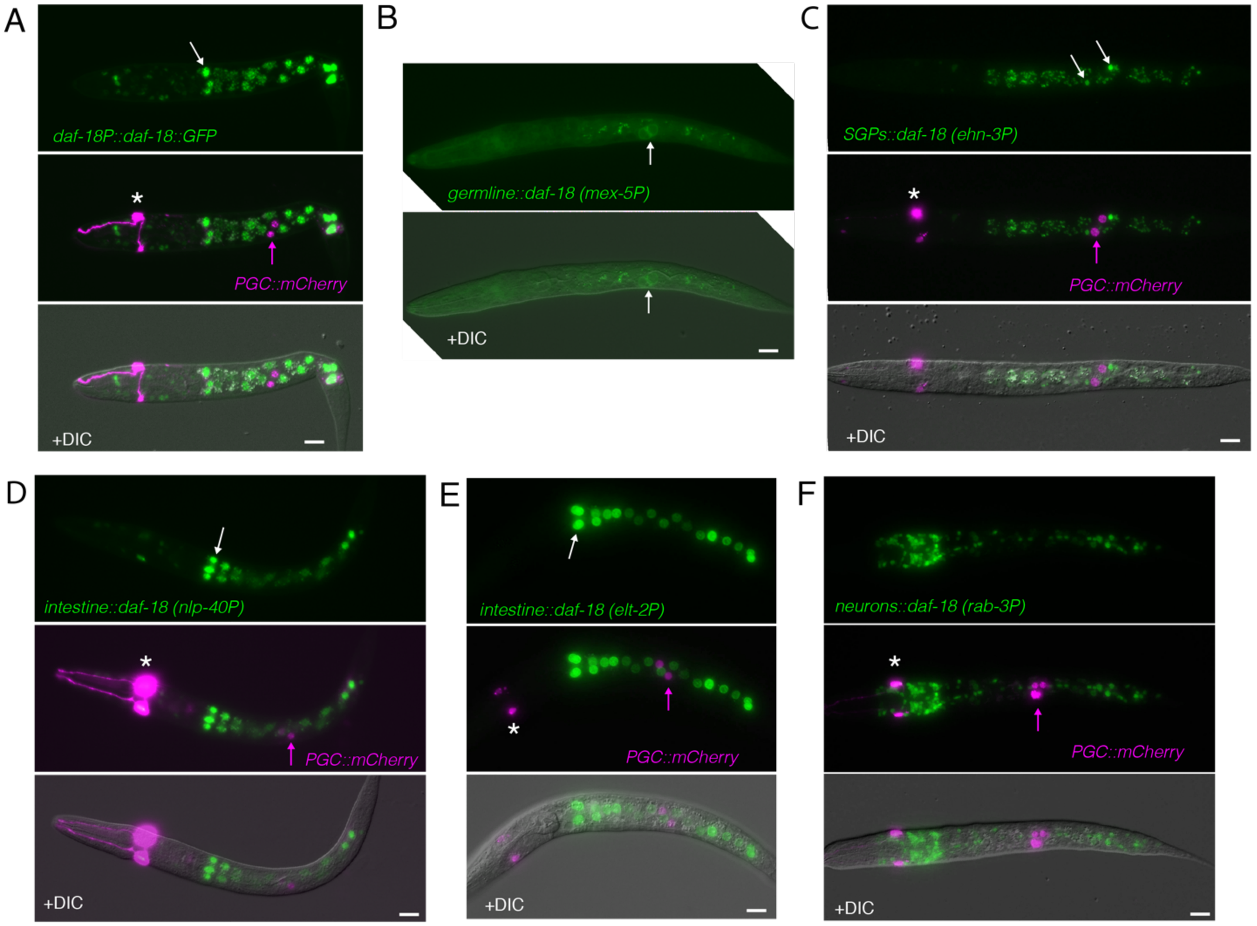


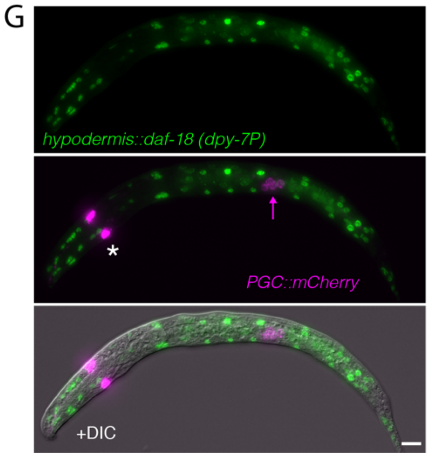


H

| **Tissue-restricted promoters** | **Germline expression of gene (TPM)*, **** | **Somatic expression of gene (TPM)*, ***** |
|---|---|---|
| *daf-18* | 90-800 | 20-300 ****, 0-20 # |
| *mex-5 (germline)* | 50-700 | 3-35 $ |
| *ehn-3 (somatic gonad precursors)* | 0-8 | 500-15500 |
| *nlp-40 (intestine)* | 1-5 | 80-500 |
| *elt-2 (intestine)* | 0 | 60-300 |
| *rab-3 (neurons)* | 0 | 50-2500 |
| *dpy-7 (hypodermis)* | 1-4 | 100-3500 |

**Figure S2. Expression patterns of *daf-18(+)::GFP* constructs.**

**A-G.** All animals are *daf-18(ok480)* mutants, and all carry *PGC::mCherry* (*naSi2*), except for panel B. Scale bars represent 10μm, white arrows indicate GFP expression in expected tissues-of-interest, pink arrows indicate PGCs, and white stars indicate a transgenic co-injection marker (*flp-17P::dsRed*) which was used to identify all extrachromosomal array-bearing animals. All *daf-18(+)::GFP* constructs are carried as extrachromosomal arrays, except for *germline::daf-18* (in panel B). Arrays carry the *daf-18* 3’UTR on the *daf-18* rescuing portion, and the *unc-54* 3’UTR for the GFP reporter portion to allow broad somatic expression (Hunt-Newbury et al. 2007) (e.g. *rab-3P::daf-18(+)::daf-18 3’UTR::SL2::GFP::H2B::unc-54 3’UTR*). **B.** The *germline:: daf-18(+)* construct (*mex-5P::SV40 NLS::daf-18 coding region::nos-2 3’UTR::SL2::GFPo::PH::tbb-2 3’UTR)* carries the *nos-2* 3’UTR on the *daf-18* rescuing portion (as used in our *PGC::mCherry*, seen in the other panels of this figure), which restricts expression to the PGCs, except for 1 additional head cell in ~10% of animals. However, the *tbb-2* 3’UTR was used for the GFP portion, which allowed broad expression. Thus, GFP is visible in somatic tissues, in addition to high expression in the germ line, but *daf-18(+)* expression should be limited to PGCs, as is the *PGC::mCherry* reporter that expresses mCherry from the same *mex-5* promoter and the same *nos-2 3’* sequences. **H.** Approximate embryonic transcript levels (Packer et al. 2019), in germline and somatic tissues of interest, of genes for which promoters were used to drive tissue-specific expression. *Adjusted transcripts per million (TPM), **Germline expression is an approximate range (from Germline:pseudotime bin 1, 2, and 3), ***Approximate range of highest-expressing cells within tissue of interest, ****Approximate range in chemosensory neurons, # Approximate range in intestine, $ Approximate range in somatic cells (neurons, body wall muscle, intestine, other).

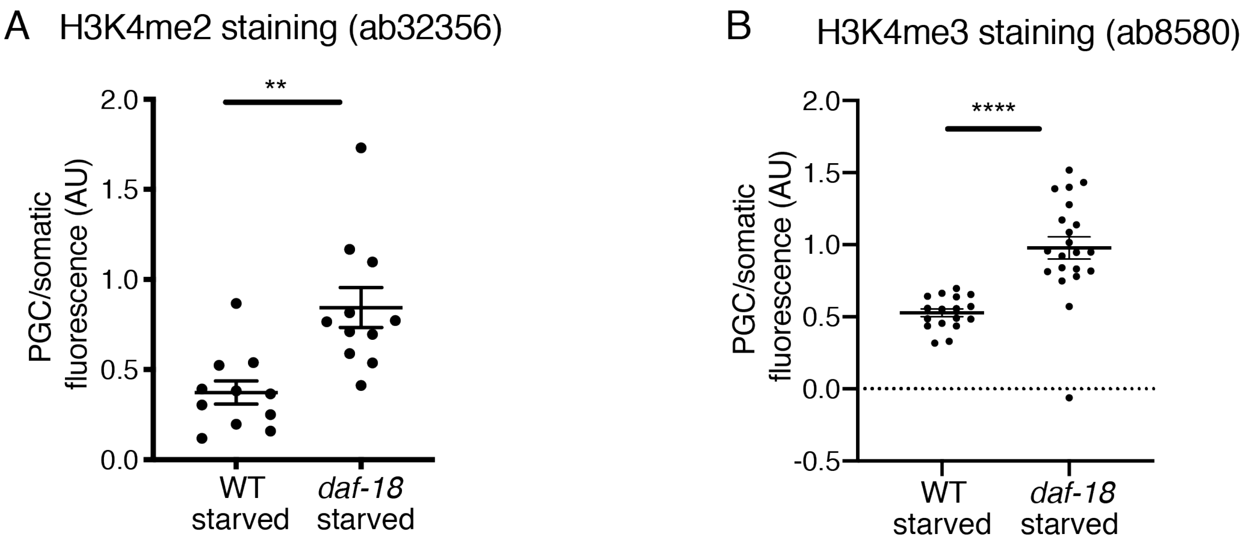


**Figure S3. Marks of active transcription are inappropriately elevated in the PGCs of starved *daf-18* mutants (additional L1 time points, and with an additional antibody).**

**A-B**. Mean staining fluorescence was measured for each PGC nucleus and its nearest somatic cell nucleus (from single image slices), and is displayed in the graph as a ratio. **A.** Antibody ab32356 (H3K4me2) was used to stain wild type (WT) and *daf-18(ok480)* mutant L1s, starved 6-8 hours after hatching (synchronized within 2 hours of hatching and imaged 6 hours later). PGCs were identified with anti-PGL-1 antibody (OIC1D4). Each dot represents a single PGC/somatic cell pair (sometimes 2 values per animal). **B.** Antibody ab8580 (H3K4me3) was used to stain WT and *daf-18(ok480)* mutant L1s hatched and starved overnight (~16h). PGCs were identified with encoded *glh-1::GFP*. Each dot represents one value per worm (2 PGC/somatic values averaged). Statistical significance determined by two-tailed T-test. $^{**}p<0.01$,$^{***}p<0.001$ $^{****}p<0.0001$.

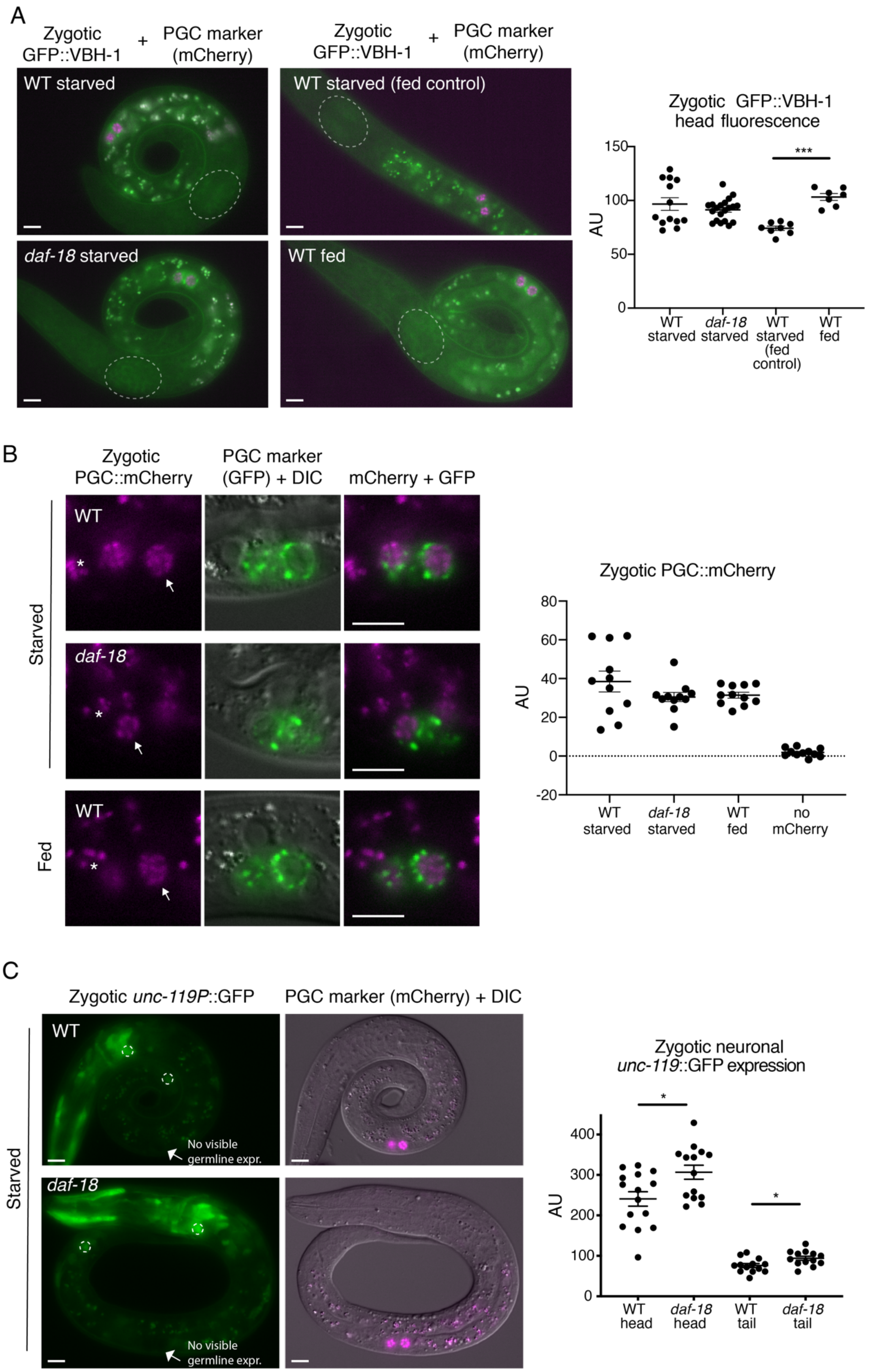
A
Zygotic GFP::VBH-1
+
PGC marker (mCherry)
WT starved
WT starved (fed control)
daf-18 starved
WT fed
Zygotic GFP::VBH-1 head fluorescence
***
150
100
50
0
AU
WT starved
daf-18 starved
WT starved (fed control)
WT fed
B
Zygotic PGC::mCherry
PGC marker (GFP) + DIC
mCherry + GFP
Starved
Fed
WT
daf-18
*
Zygotic PGC::mCherry
80
60
40
20
0
-20
AU
WT starved
daf-18 starved
WT fed
no mCherry
C
Zygotic unc-119P::GFP
PGC marker (mCherry) + DIC
Starved
WT
daf-18
No visible germline expr.
Zygotic neuronal unc-119::GFP expression
*
400
300
200
100
0
AU
WT head
daf-18 head
WT tail
daf-18 tail

**Figure S4. Further zygotic transgene expression analysis.**

**A-C.** Images show epifluorescence. Mean fluorescence intensity (AU=Arbitrary Units) values are plotted to the right of each set of images. Each dot represents one animal. Non-cross progeny were identified by the lack of somatic GFP (A and C) or lack of PGC::mCherry (B), which are all visible when present, even in starved animals. Mean +/- SEM. All scale bars represent 10μm. **A.** Zygotic GFP::VBH-1 expression in the soma (head neurons) is not expressed more highly in starved *daf-18* mutants than starved wild type (WT). Dashed ellipses show regions measured for mean fluorescence. "Starved" animals were imaged within 5 hours of hatching (no food). "Starved (fed control)" animals were allowed to hatch for up to 22h after clean embryo preparation, and "fed" animals were fed for 5 hours after being allowed to hatch for up to 17h, and both sets were imaged around the same time. Graph shows the pooled results of 2 experiments. Statistical significance determined by one-way ANOVA with Tukey's multiple comparisons test. ***$p<0.001$. **B.** Zygotic nuclear PGC::mCherry expression is not expressed more highly in starved *daf-18* mutant PGCs than starved WT PGCs. Arrows point to zygotic PGC::mCherry expression in PGCs, and asterisks mark auto-fluorescent gut granules. Nuclear PGC::mCherry fluorescence was measured by hand-drawing regions of interest around PGC nuclei. One experimental replicate is graphed, and 2 others showed similar results (data not shown). Statistical significance determined by one-way ANOVA with Tukey's multiple comparisons test. **C.** Zygotic *unc-119P*::GFP is not expressed in the PGCs of starved *daf-18* mutants. However, it is expressed at a slightly higher level in neurons of starved *daf-18* mutants. L1s were imaged up to 18 hours after clean embryo preparation. Dashed circles indicate approximate neuronal regions measured (one head neuron and one tail neuron per animal). Graph shows the pooled results of 2 experiments. Statistical significance determined by two-tailed T-test. *$p<0.05$.

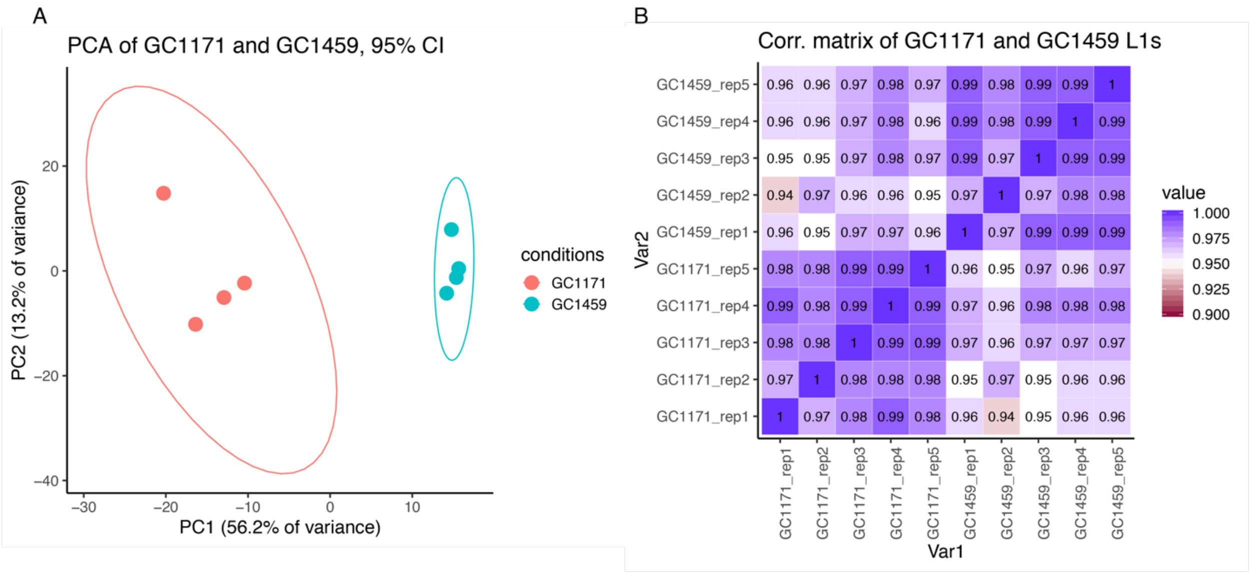


**Figure S5-1. RNA Sequencing: quality control analyses.**

**A.** Principal component analysis (PCA) of GC1171 (wild type) and GC1459 (*daf-18(ok480)*) starved L1s in RNA Sequencing experiment, highlighting low variance (13.2%) within each genotype, and high variance (56.2%) between genotypes. **B.** Correlation matrix of GC1171 (wild type) and GC1459 (*daf-18(ok480)*) showing that replicates of the same genotype were highly correlated.

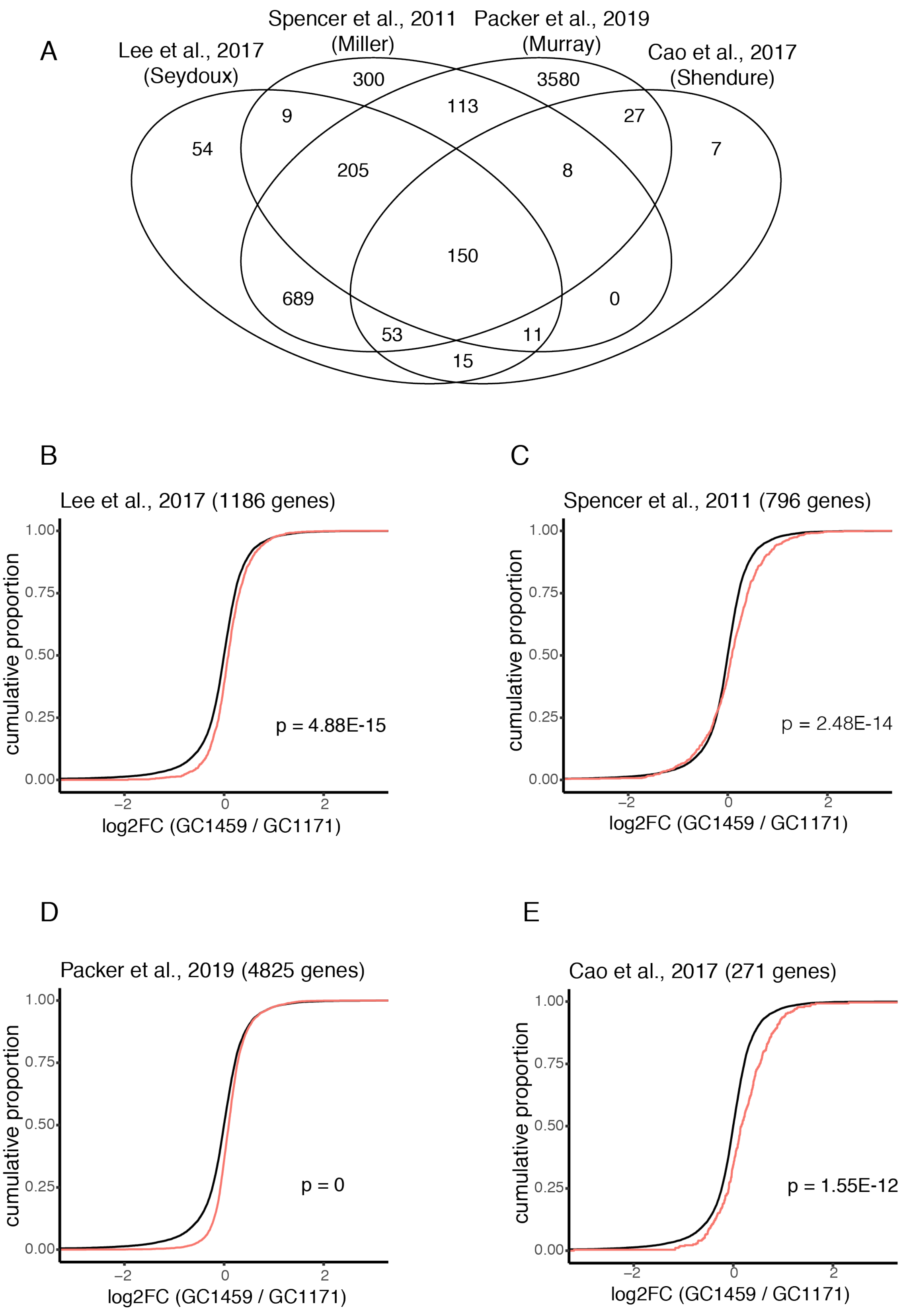

A
Lee et al., 2017 (Seydoux)
Spencer et al., 2011 (Miller)
Packer et al., 2019 (Murray)
Cao et al., 2017 (Shendure)
54
9
300
113
3580
27
7
205
8
150
689
0
53
11
15
B
Lee et al., 2017 (1186 genes)
cumulative proportion
p = 4.88E-15
log2FC (GC1459 / GC1171)
C
Spencer et al., 2011 (796 genes)
cumulative proportion
p = 2.48E-14
log2FC (GC1459 / GC1171)
D
Packer et al., 2019 (4825 genes)
cumulative proportion
p = 0
log2FC (GC1459 / GC1171)
E
Cao et al., 2017 (271 genes)
cumulative proportion
p = 1.55E-12
log2FC (GC1459 / GC1171)

**S5-2. RNA sequencing further germline gene analyses**

**A.** Venn diagram showing the overlap of all four germline gene sets analyzed (Cao et al. 2017; Lee et al. 2017; Packer et al. 2019; Spencer et al. 2011), and numbers of genes from these sets detected in our RNA Sequencing experiment. **B-E.** Cumulative distribution function (CDF) plots for individual published germline gene sets (red lines) compared to the background set of all transcripts detected in our experiment (12,592 genes). p-values determined by Kolmogorov-Smirnov (KS) test are shown. Numbers of transcripts from published gene sets detected in our experiment are shown in graph titles (parentheses).

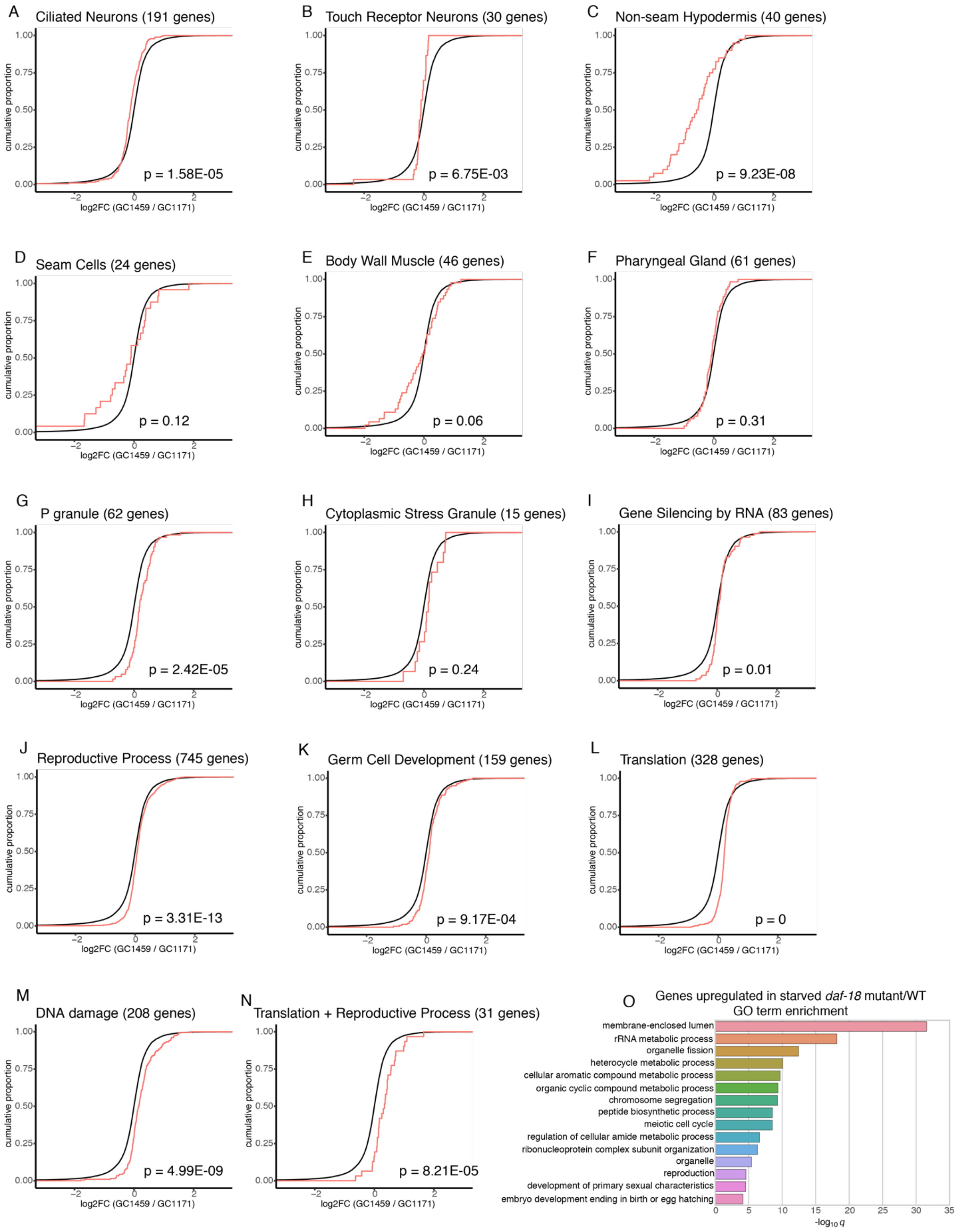
A
Ciliated Neurons (191 genes)
p = 1.58E-05
B
Touch Receptor Neurons (30 genes)
p = 6.75E-03
C
Non-seam Hypodermis (40 genes)
p = 9.23E-08
D
Seam Cells (24 genes)
p = 0.12
E
Body Wall Muscle (46 genes)
p = 0.06
F
Pharyngeal Gland (61 genes)
p = 0.31
G
P granule (62 genes)
p = 2.42E-05
H
Cytoplasmic Stress Granule (15 genes)
p = 0.24
I
Gene Silencing by RNA (83 genes)
p = 0.01
J
Reproductive Process (745 genes)
p = 3.31E-13
K
Germ Cell Development (159 genes)
p = 9.17E-04
L
Translation (328 genes)
p = 0
M
DNA damage (208 genes)
p = 4.99E-09
N
Translation + Reproductive Process (31 genes)
p = 8.21E-05
cumulative proportion
0.00
0.25
0.50
0.75
1.00
−2
0
2
log2FC (GC1459 / GC1171)
O
Genes upregulated in starved daf-18 mutant/WT
GO term enrichment
membrane-enclosed lumen
rRNA metabolic process
organelle fission
heterocycle metabolic process
cellular aromatic compound metabolic process
organic cyclic compound metabolic process
chromosome segregation
peptide biosynthetic process
meiotic cell cycle
regulation of cellular amide metabolic process
ribonucleoprotein complex subunit organization
organelle
reproduction
development of primary sexual characteristics
embryo development ending in birth or egg hatching
0
5
10
15
20
25
30
35
-log10 q

**S5-3. RNA Sequencing: analyses of tissues and GO terms**

Cumulative distribution function (CDF) plots of transcript sets of interest (red line) versus the background set of all 12,592 transcripts detected (black line). p-values determined by Kolmogorov-Smirnov (KS) test are shown. Numbers of transcripts from published gene sets detected in our experiment are shown in graph titles (parentheses). **A-F.** CDF plots for lists of transcripts from tissues of interest known to express *daf-18* (ciliated neurons, non-seam hypodermis, seam cells, body wall muscle), and tissues that may not express *daf-18* (touch receptor neurons, pharyngeal gland) (Cao et al. 2017; Packer et al. 2019). Tissue-specific gene lists were generated using the GExplore1.4 web tool (underlying data from Cao et al., 2017). **G-N.** CDF plots for lists of transcripts associated with specific gene ontology (GO) terms (using WormBase). **O.** Unbiased analysis of gene ontology (GO) terms, using WormBase's Enrichment Analysis Tool.
