## Supplementary material for "DAF-18/PTEN inhibits germline zygotic gene activation during primordial germ cell quiescence": Table S1

**Table S1. Strains, plasmids and primers used in this study**

**Strains used in this study**

| **Strain** | ***Genotype*** | **Construction details/Source** | **Source/Reference** | **Abbreviation in text** |
|---|---|---|---|---|
| N2 | *Wild-type* | | (Brenner 1974) | |
| GC1171 | *naSi2 (mex-5p::mCherry::H2B::nos-2 3'UTR +unc-119(+)) II; unc-119(ed3)? III* | *naSi2 [pGC550 (mex-5p::mCherry::H2B::nos-2 3'UTR +unc-119(+))]* | (Roy et al. 2018) | "PGC::mCherry", "WT" |
| GC1459 | *naSi2 (mex-5p::mCherry::H2B::nos-2 3'UTR +unc-119(+)) II; unc-119(ed3) III ?; daf-18(ok480) IV* | | this work | "PGC::mCherry; daf-18" |
| DUP64 | *glh-1(sam24[glh-1::gfp::3xFLAG]) I* | Gift from Dustin Updike | (Andralojc et al. 2017) | *"glh-1::GFP,"* "WT" |
| GC1527 | *glh-1(sam24[glh-1::gfp::3xFLAG]) I; daf-18(ok480) IV* | | this work | *"glh-1::GFP; daf-18"* |
| FT95 | *unc-119(ed3) III; xnIs23 [cdc-42P::ZF1-GFP-CDC-42, unc-119(+)]* | Gift from Jeremy Nance | (Armenti et al. 2014) | |
| OH441 | *otIs45 [unc-119P::GFP]* | CGC | (Altun-Gultekin et al. 2001) | |
| GC1537 | *vbh-1(na110[GFP::vbh-1]) I; naSi2 (mex-5p::mCherry::H2B::nos-2 3'UTR +unc-119(+)) II; unc-119(ed3)? III* | ***na110[GFP::vbh-1]*** : Inserted GFP into N terminus of endogenous vbh-1 gene using CRISPR-based "hybrid" GFP donor method. Injected into GC1171 (naSi2): Cas9 (IDT) at .25ug/uL, tracrRNA (IDT) at 0.1 ug/uL, vbh-1 target crRNA at 0.056 ug/uL, dpy-10 crRNA (IDT) at 0.0056 ug/uL, dpy-10 ss oligo at 0.4 uM, sticky repair PCR (GFPo amplified from pGC689 with 120bp vbh-1 homology arms) at 200 ng/uL. | this work | *GFP::vbh-1* |
| GC1576 | *vbh-1(na110[GFP::vbh-1]) I; naSi2 (mex-5p::mCherry::H2B::nos-2 3'UTR +unc-119(+)) II; unc-119(ed3)? III; daf-18(ok480) IV* | | this work | |
| HT1593 | *unc-119(ed3) III* | CGC | Heidi Tissenbaum via CGC | |
| GC1456 | *naSi2 (mex-5p::mCherry::H2B::nos-2 3'UTR +unc-119(+)) II; unc-119(ed3) III ?; daf-18(ok480) IV; naEx261 (daf-18P::daf-18::daf-18 3'UTR::SL2::GFP::H2B::unc-54 3'UTR)* | **daf-18 genomic "rescue," *naEx261*** = Injected into GC1459 (*naSi2; daf-18* ) : *pGC688 [daf-18P::daf-18::daf-18 3'UTR::SL2::GFP::H2B::unc-54 3'UTR]* at 50ng/uL, pJB134 [*flp-17P::dsRed* ] at 50ng/uL, 1kb DNA ladder at 25ng/uL. | this work | |
| GC1460 | *naSi2 (mex-5p::mCherry::H2B::nos-2 3'UTR +unc-119(+)) II; unc-119(ed3) III ?; daf-18(ok480) IV; naEx264 (rab-3P::daf-18::daf-18 3'UTR::SL2::GFP::H2B::unc-54 3'UTR)* | **daf-18 neuronal "rescue," *naEx264*** = Injected into GC1459 (*naSi2; daf-18* ) : pGC694 [*rab-3P::daf-18::daf-18 3'UTR::SL2::GFP::H2B::unc-54 3'UTR* ] at 20 ng/uL, pJB134 [*flp-17P::dsRed* ] at 50ng/uL, 1kb DNA ladder at 60ng/uL. | this work | |
| GC1464 | *naSi2 (mex-5p::mCherry::H2B::nos-2 3'UTR +unc-119(+)) II; unc-119(ed3) III ?; daf-18(ok480) IV; naEx268 (nlp-40P::daf-18::daf-18 3'UTR::SL2::GFP::H2B::unc-54 3'UTR)* | **daf-18 intestinal "rescue," *naEx268*** = Injected into GC1459 (*naSi2; daf-18* ) : pGC695 [*nlp-40P::daf-18::daf-18 3'UTR::SL2::GFP::H2B::unc-54 3'UTR* ] at 50 ng/uL, pJB134 [*flp-17P::dsRed* ] at 50ng/uL, 1kb DNA ladder at 25ng/uL. | this work | |
| GC1465 | *naSi2 (mex-5p::mCherry::H2B::nos-2 3'UTR +unc-119(+)) II; unc-119(ed3) III ?; daf-18(ok480) IV; naEx269 (dpy-7P::daf-18::daf-18 3'UTR::SL2::GFP::H2B::unc-54 3'UTR)* | **daf-18 hypodermal "rescue," *naEx269*** = Injected into GC1459 (*naSi2; daf-18* ) : pGC696 [*dpy-7P::daf-18::daf-18 3'UTR::SL2::GFP::H2B::unc-54 3'UTR* ] at 10ng/uL, pJB134 [*flp-17P::dsRed* ] at 50ng/uL, 1kb DNA ladder at 60ng/uL. | this work | |
| GC1485 | *naSi2 (mex-5p::mCherry::H2B::nos-2 3'UTR +unc-119(+)) II; unc-119(ed3) III ?; daf-18(ok480) IV; naEx275 (elt-2P::daf-18::daf-18 3'UTR::SL2::GFP::H2B::unc-54 3'UTR)* | **daf-18 intestinal "rescue," *naEx275*** = Injected into GC1459 (*naSi2; daf-18* ) : pGC716 [*elt-2P::daf-18::daf-18 3'UTR::SL2::GFP::H2B::unc-54 3'UTR* ] at 20ng/uL, pJB134 [*flp-17P::dsRed* ] at 50ng/uL, 1kb DNA ladder at 60ng/uL. | this work | |

| | | | |
|---|---|---|---|
| GC1484 | *naSi2 (mex-5p::mCherry::H2B::nos-2 3'UTR +unc-119(+)) II; unc-119(ed3) III ?; daf-18(ok480) IV; naEx274 (ehn-3P::daf-18::daf-18 3'UTR::SL2::GFP::H2B::unc-54 3'UTR)* | **daf-18 somatic gonad precursor (SGP) "rescue," *naEx274*** = Injected into GC1459 (*naSi2; daf-18* ) : pGC717 [*ehn-3P::daf-18::daf-18 3'UTR::SL2::GFP::H2B::unc-54 3'UTR* ] at 20ng/uL, pJB134 [*flp-17P::dsRed* ] at 50ng/uL, 1kb DNA ladder at 60ng/uL. | this work |
| GC1479 | *naSi22 (mex-5P::daf-18::nos-2 3'UTR::SL2::GFPo::PH::tbb-2 3'UTR + unc-119(+)) II; unc-119(ed3) III ?; daf-18(ok480) IV* | **daf-18 germline "rescue," *naSi22*** = Used Cas9-triggered long-range HDR, injected into strain HT1593 (*unc-119(ed3) III)* : pGC689 *[mex-5P::daf-18::nos-2 3'UTR::SL2::GFPo::PH::tbb-2 3'UTR +unc-119(+)* ] at 20ng/uL, pDD122 at 50ng/uL, pMA122 at 10ng/uL, pGH8 at 10 ng/uL, pCFJ90 at 2.5 ng/uL, pCFJ104 at 5 ng/uL, 1kb DNA ladder at 20ng/uL. | this work |
| GC1483 | *glh-1(sam24[glh-1::gfp::3xFLAG]) I; naSi22 (mex-5P::daf-18::nos-2 3'UTR::SL2::GFPo::PH::tbb-2 3'UTR) II; unc-119(ed3) III ?; daf-18(ok480) IV* | | this work |

**Plasmids used in this study**

| Plasmid name | Description | Reference/Details/Construction | Source/Reference |
|---|---|---|---|
| pJB134 | *flp-17P::dsRed* (coinjection marker) | | Gift from N. Ringstad (Julia Brandt) |
| pGC550 | Source of *H2B* , *nos-2* 3'UTR, [*mex-5P::FRT::mCherry::H2B::nos-2 3'UTR::FRT::GFP::H2B* ] | | (Roy et al. 2018) |
| pEM1 | Source of *SL2::GFP::unc-54 3' UTR* | | (Macosko et al. 2009) |
| pGC658 | Source of ttTi5605 left and right homology arms, C. briggsae unc-119(+), *mex-5P* , and *SL2::GFPo::PH::tbb-2 3'UTR* | | this study |
| pBD001 | Source of *elt-2P* | | Gift from J. Nance (Brielle Dojer) |
| pYA12 | Source of *ehn-3P* | | Gift from J. Nance (Yusuff Abdu) |
| pGC688 | *daf-18* genomic "rescue" construct, [*daf-18P::daf-18::daf-18 3'UTR::SL2::GFP::H2B::unc-54 3'UTR* ] | Gibson Assembly used to combine: *daf-18* full promoter to the next gene start (~800bp)::*daf-18* coding region with introns (4723bp)::*daf-18* 3'UTR (439bp) [Amplified from genome], *SL2::GFP* [pEM1], *H2B* [pGC550], *unc-54* 3'UTR [pEM1]. Note: pGC688 carries a single nucleotide insertion in the promoter. | this study |
| pGC689 | *daf-18* germline "rescue" construct, [*mex-5P::daf-18::nos-2 3'UTR::SL2::GFPo::PH::tbb-2 3'UTR* ] | Gibson Assembly used to combine: ttTi5605 left homology arm::C. briggsae *unc-119(+)::mex-5P* (487bp) [pGC658], *daf-18* coding region with introns (4723bp) [pGC688], *nos-2* 3'UTR (836bp) [pGC550], *SL2::GFPo* (codon-optimized)::*PH::tbb-2* (332bp) 3'UTR::ttTi5605 right homology arm [pGC658] | this study |
| pGC694 | *daf-18* neuronal "rescue" construct, [*rab-3P::daf-18::daf-18 3'UTR::SL2::GFP::H2B::unc-54 3'UTR* ] | Gibson Assembly used to combine: *rab-3P* (1208 bp) [amplified from genome], *daf-18* coding region with introns (4723bp)::*daf-18* 3'UTR (439bp)::*SL2::GFP::H2B::unc-54* 3'UTR [pGC688]. *rab-3* promoter as used in {Wang, 2017 #186}. | this study |
| pGC695 | *daf-18* intestinal "rescue" construct 1, [*nlp-40P::daf-18::daf-18 3'UTR::SL2::GFP::H2B::unc-54 3'UTR* ] | Gibson Assembly used to combine: *nlp-40P* (3511 bp) [amplified from genome], *daf-18* coding region with introns (4723bp)::*daf-18* 3'UTR (439bp)::*SL2::GFP::H2B::unc-54* 3'UTR [pGC688]. *nlp-40* promoter as used in {Wang, 2017 #186}. | this study |

| | | | |
|---|---|---|---|
| pGC696 | *daf-18* hypodermal "rescue" construct, [*dpy-7P::::daf-18::daf-18 3'UTR::SL2::GFP::H2B::unc-54 3'UTR* ] | Gibson Assembly used to combine: *dpy-7P* (339 bp) [amplified from genome], *daf-18* coding region with introns (4723bp)::*daf-18 3'UTR (439bp)::SL2::GFP::H2B::unc-54 3'UTR* [pGC688]. *dpy-7* promoter as used in {Wang, 2017 #186}. | this study |
| pGC716 | *daf-18* intestinal "rescue" construct 2, [*elt-2P::daf-18::daf-18 3'UTR::SL2::GFP::H2B::unc-54 3'UTR* ] | Gibson Assembly used to combine: *elt-2P* (3917 bp) [pBD001], *daf-18* coding region with introns (4723bp)::*daf-18* 3'UTR (439bp)::*SL2::GFP::H2B::unc-54* 3'UTR [pGC688]. | this study |
| pGC717 | *daf-18* somatic gonad precursor (SGP) "rescue" construct , [*ehn-3::daf-18::daf-18 3'UTR::SL2::GFP::H2B::unc-54 3'UTR* ] | Gibson Assembly used to combine: *ehn-3P* (2792 bp) [pYA12], *daf-18* coding region with introns (4723bp)::*daf-18* 3'UTR (439bp)::*SL2::GFP::H2B::unc-54* 3'UTR [pGC688]. | this study |
| pDD122 | Cas9 + sgRNA | *Peft-3::Cas9* + ttTi5605 sgRNA. Plasmid that is targeted to a genomic site near the ttTi5605 Mos1 insertion allele . | (Dickinson et al. 2013) |
| pMA122 | *Phsp::peel-1* | Negative selection marker | (Frokjaer-Jensen et al. 2012) |
| pGH8 | *Prab-3::mCherry* (Pan-neuronal) | | (Kihira et al. 2012) |
| pCFJ90 | *Pmyo-2::mCherry* (pharynx muscle) | | (Frokjaer-Jensen et al. 2008) |
| pCFJ104 | *Pmyo-3::mCherry* (body wall muscle) | | (Frokjaer-Jensen et al. 2008) |

**Primers used in this study**

| Primer name | Description | Sequence | Reference |
|---|---|---|---|
| *rab-3P* into pGC688 FWD | Gibson Assembly Primer. Capital letter nucleotides are corresponds to pGC688, lowercase corresponds to rab-3P | TTTGCTGGCCTTTTGCTCACgatcttcagatgggagcagt | (Wang et al. 2017) |
| *rab-3P* into pGC688 REV | Gibson Assembly Primer. Capital letter nucleotides are corresponds to pGC688, lowercase corresponds to rab-3P | TCTGGAGGAGGAGTAACCATctgaaaatagggctactgtaga | (Wang et al. 2017) |
| *nlp-40P* into pGC688 FWD | Gibson Assembly Primer. Capital letter nucleotides are corresponds to pGC688, lowercase corresponds to nlp-40P | TTTGCTGGCCTTTTGCTCACgtgagaagatatcgtacgagg | (Wang et al. 2017) |
| *nlp-40P* into pGC688 REV | Gibson Assembly Primer. Capital letter nucleotides are corresponds to pGC688, lowercase corresponds to nlp-40P | TCTGGAGGAGGAGTAACCATgttgattgtgtatgtttggct | (Wang et al. 2017) |
| *dpy-7P* into pGC688 FWD | Gibson Assembly Primer. Capital letter nucleotides are corresponds to pGC688, lowercase corresponds to dpy-7P | TTTGCTGGCCTTTTGCTCACtcattccacgatttctcgca | (Wang et al. 2017) |
| *dpy-7P* into pGC688 REV | Gibson Assembly Primer. Capital letter nucleotides are corresponds to pGC688, lowercase corresponds to dpy-7P | TCTGGAGGAGGAGTAACCATtctggaacaaaatgtaagaatattct | (Wang et al. 2017) |
| *elt-2P* into pGC688 FWD | Gibson Assembly Primer. Capital letter nucleotides are corresponds to pGC688, lowercase corresponds to elt-2P | TTTGCTGGCCTTTTGCTCACgatcttctccttcccatgtgctg | this work |
| *elt-2P* into pGC688 REV | Gibson Assembly Primer. Capital letter nucleotides are corresponds to pGC688, lowercase corresponds to elt-2P | TCTGGAGGAGGAGTAACCATtctataatctattttctagtttctattttattagaatgcc | this work |
| *ehn-3P* into pGC688 FWD | Gibson Assembly Primer. Capital letter nucleotides are corresponds to pGC688, lowercase corresponds to ehn-3P | TTTGCTGGCCTTTTGCTCACGaattttaaatgggtattcggatgtaaaatgatgagaaag | this work |

| | | | |
|---|---|---|---|
| *ehn-3P* into pGC688 REV | Gibson Assembly Primer. Capital letter nucleotides are corresponds to pGC688, lowercase corresponds to ehn-3P | TCTGGAGGAGGAGTAACCATtttgtaatttggaagctgggag | this work |
| GCo1207 | *daf-18(ok480)* detection FWD | gagtcggtggtccatttgagatacc | this work |
| GCo1208 | *daf-18(ok480)* detection REV. GCo1207 + 1208 = 1166 bp in WT, 211 bp in *daf-18 (ok480)* | actggcaacgaatgaatacgcaggt | this work |
| GCo2250 | *daf-18(ok480)* detection REV in deletion. GCo1207+GCo2250 = 775 bp in WT, no band in *daf-18(ok480)* | GCAGGAGGAAGTTCGTCTTCAG | this work |
| *vbh-1* crRNA | | CATtgtaagactatatcaaa | this work |
| *vbh-1* /GFPo sticky repair template PCR FWD | to insert GFPo (codon-optimized) into vbh-1 N terminus. Lower case is vbh-1 locus homology, upper case is GFPo | taaaatgaatttcagatacatattctgtaatctcgacgtcaattgggtacaaaatcggtttagattcgccgatttttctttttcgaattccgggtttccgtttgatatagtcttacaATGAGTAAAGGAGAAGAATTGTT | this work |
| *vbh-1* /GFPo sticky repair template PCR FWD | to insert GFPo (codon-optimized) into vbh-1 N terminus. Lower case is vbh-1 locus homology, upper case is GFPo | gatgaccagctgaaaaaatgaggttgaaatcttgaaatccgtttcgaaatcatacccttgtccctgttgttgttgtggcggctgttgttgaccgttgtgattcgcataatattgtgtgttCTTGTAGAGCTCGTCCATTC | this work |
| *dpy-10* ssDNA oligo | for *dpy-10* co-CRISPR | CACTTGAACTTCAATACGGCAAGATGAGAATGACTGGAAACCGTACCGCATGCGGTGCCTATGGTAGCGGAGCTTCACATGGCTTCAGACCAACAGCCTAT | (Paix et al. 2014) |
| *dpy-10* crRNA | for *dpy-10* co-CRISPR | GCUACCAUAGGCACCACGAG | (Paix et al. 2014) |
| GCo2332 | fragment 1 FWD for daf-18 genomic array. Capital letter nucleotides correspond to the vector and lowercase correspond to *daf-18* locus | TTTGCTGGCCTTTTGCTCACatttcatcgcactatacaaactttagactctaaataca | this work |
| GCo2334 | fragment 1 REV for *daf-18* genomic array. | ttgaattcaaaaagaaaaataaattggagccaacga | this work |
| GCo2335 | fragment 2 FWD for *daf-18* genomic array. | cacttcgttggctccaatttatttttctttttga | this work |
| GCo2336 | fragment 2 REV for *daf-18* genomic array. Capital letter nucleotides correspond to the vector and lowercase correspond to *daf-18* locus | TGAAAGTAGGATGAGACAGCcgaatgcgcctagattctatgtatcg | this work |
